# European common frog (*Rana temporaria*) recolonised Switzerland from multiple glacial refugia in northern Italy via trans- and circum-Alpine routes

**DOI:** 10.1101/696153

**Authors:** A Jansen van Rensburg, M Robin, B C Phillips, J Van Buskirk

## Abstract

The high mountain ranges of western Europe have had a profound effect on the recolonisation of Europe from glacial refugia. The Alps present a particularly interesting case, because they present an absolute barrier to dispersal to most lineages, obstructing recolonisation from multiple refugia in the Italian Alps. Here we investigate the effect of the European Alps on the the phylogeographic history of *Rana temporaria* across its range in Switzerland. Based on partial *cytochrome b* and *COX1* sequences we find two mitochondrial lineages that occur roughly north and south of the alpine ridge bisecting Switzerland, with contact zones between them in the east and west. The northern haplogroup falls within the previously identified common western European haplogroup, while the southern haplogroup is unique to Switzerland. We find that the lineages diverged ~110 kya, approximately the onset of the last glacial maximum, indicative of origins in separate refugia. Phylogenetic analyses suggest that the lineages originate from two refugia in northern Italy, and colonised Switzerland via trans- and curcum-alpine routes. Our results show that the European Alps is a semi-permeable barrier to dispersal for *R. temporaria*, and have contributed to the complex recolonisation history of Switzerland.

## Introduction

Contemporary geographic distributions of genetic variation result from processes occurring on diverse time scales. Climate oscillations and associated ice-ages that occurred during the last ~700 ky had a major influence on contemporary European ecosystems (Taberlet *et al.* 1998; Hewitt 1999, 2000). The extent of the ice-sheets and inhospitable habitat restricted many species to three major refugia: the Balkans, Italy, and the Iberian Peninsula (Taberlet *et al.* 1998). After the retreat of the ice-sheets, Europe was recolonised by the surviving populations. Differing dispersal abilities, residual habitat discontinuities, and geographic complexities have determined these recolonisation routes. This has had a fundamental impact on the extant geographic distribution of species and their genetic diversity (Roy *et al.* 1996; Taberlet *et al.* 1998; Hewitt 2000; Willis & Whittaker 2008; Avise & Riddle 2009).

High mountain ranges such as the European Alps and Pyrenees were the last to deglaciate. For taxa adapted to warm climates, the persistence and extent of alpine ice-sheets and the east-west orientation of the mountain ranges hindered biotic recolonisation of Europe from their refugia (Taberlet *et al.* 1998). Cold-tolerant organisms exhibit a more complex history. Many expanded their range during glacial periods, some persisted in nunataks within the Alps, and trans-alpine recolonisation has occurred in several species (Mátyás & Sperisen 2001; Lugon-Moulin & Hausser 2002; Parisod 2008; Yannic *et al.* 2008; Braaker & Heckel 2009). However, the effect of the Alps on the biogeography of cold-tolerant species remains understudied, particularly for vertebrates (but see Yannic *et al.* 2008, Braaker & Heckel 2009).

Here we describe the phylogeographic history of a cold-tolerant amphibian, the European common frog (*Rana temporaria*) by densely sampling populations across the Alps in Switzerland. This species is the most widespread anuran in Europe, occurring from northern Italy and Spain to the sub-arctic tundra in Fennoscandia in the north and the Ural mountains in the east. *R. temporaria* is ubiquitous in the European Alps and occurs up to an elevation of 2600 m (Sillero *et al.* 2014). Two deeply diverged (~0.7 Mya) mitochondrial lineages occur in western and eastern/northern Europe, respectively (Palo *et al.* 2004). A contact zone has been identified that extends from northern Germany (Schmeller *et al.* 2008) to the northern lowlands of Switzerland and north western Italy (Teacher *et al.* 2009). The precise location and structure of the contact zone in the Alps has not been resolved by existing studies (Stefani *et al.* 2012; Rodrigues *et al.* 2013). The western haplogroup has higher genetic diversity than the eastern haplogroup which suggests that *R. temporaria* recolonised western Europe from multiple glacial refugia. Both Iberia (Teacher *et al.* 2009) and Italy (Stefani *et al.* 2012) have been considered as the main refugia for the western haplogroup. However, sampling has not been available to reconstruct the main routes of post-glacial recolonisation of the western haplogroups, and it remains unclear whether the Alps was a barrier to re-colonisation into western Europe.

Our main aims are to determine the colonisation history and extant population structure of *Rana temporaria* in Switzerland. Specifically we ask whether 1) a contact zone exists between Eastern and Western haplotypes in Switzerland, 2) Switzerland was initially colonised from one or multiple glacial refugia, and 3) determine whether the Alps was a barrier to colonisation.

The geography of the Alps between northern Italy and Switzerland are characterised by two main high mountain ridges running roughly east to west. We refer to the first as the southern Alpine ridge. This comprises the highest peaks and roughly constitutes the border between Italy and Switzerland. The second ridge is called the northern Alpine ridge, and roughly bisects Switzerland into a northern and southern part.

## Materials and Methods

### Sampling

To investigate the geographic distribution of genomic variation in Switzerland, we sampled 82 populations across a ~2300 m gradient of elevation in the Swiss Alps during the 2013 breeding season (Fig. 1, Table 1). Approximately 10 eggs from each of 20-30 freshly laid clutches were collected from each site. The eggs were transported to the University of Zürich, where they were hatched in separate water-filled containers in a water bath. The tadpoles were euthanised and stored in ethanol when they reached stage 36 (Gosner, 1960). Each clutch is assumed to represent a single family; thus one tadpole per clutch was used for all work presented here.

**Figure 1.**
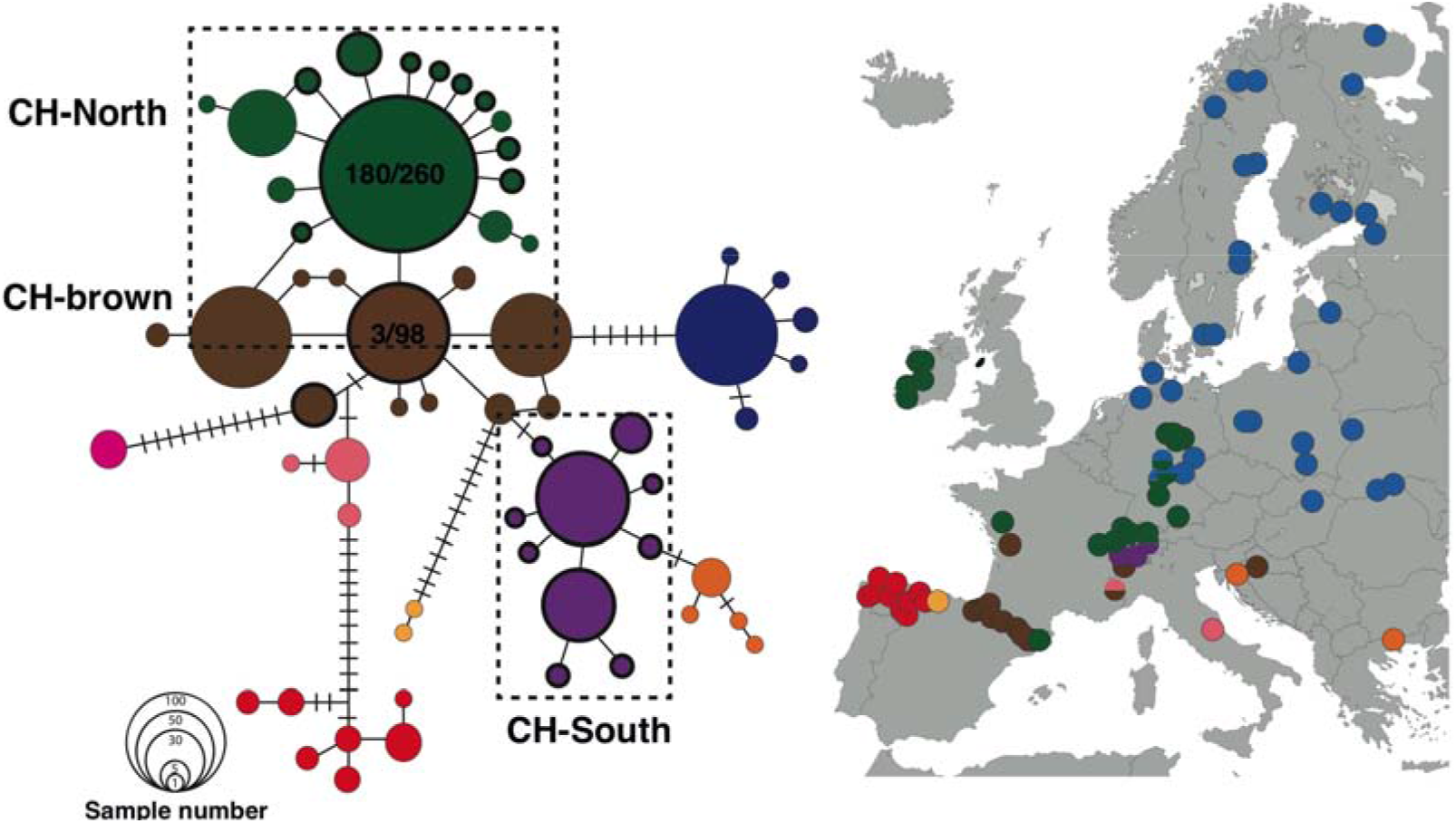
Haplotype network of 331bp of the *cytb* gene sequenced for this study and combined with data from Vences *et al.* (2013). Geographic distribution of the haplotypes within the eastern (blue) and western (other colours) clades is shown on a map of Europe. Swiss haplotypes (shown with bold outlines) fall within the green (CH-North) and brown (CH-brown) haplogroups, and a Swiss-specific purple haplogroup (CH-South). Numbers in the green and brown haplotypes show the proportion of samples from this study assigning to that haplotype.

**Table 1.** Diversity measures and neutrality tests for the *cytb*, COX1, and concatenated datasets generated in this study. CH-North includes all samples with haplotypes from the Northern haplogroup, and CH-South from the Swiss-specific haplogroup. N = Number of samples; length = sequence length in base pairs; nr Hap = the number of haplotypes detected, Hd = haplotype diversity (standard deviation); Nd = nucleotide diversity (standard deviation). Three tests for selection are included. * = *P* < 0.05; ** = *P* < 0.01; *** = *P* < 0.001.

| Pop Group | N | length (bp) | nr Hap | Hd (SD) | $\pi$ (SD) | Tajima's D | Fu & Li's D | Fu & Li's F |
| --- | --- | --- | --- | --- | --- | --- | --- | --- |
| ***cytb*** |  |  |  |  |  |  |  |  |
| All | 368 | 448 | 29 | 0.712 (0.020) | 0.00857<br>(0.00019) | -0.377 | -2.896* | -2.202 |
| CH-North | 217 | 448 | 17 | 0.347 (0.042) | 0.00126<br>(0.00021) | -2.207** | -3.945** | -3.918** |
| CH-South | 151 | 448 | 12 | 0.633 (0.034) | 0.00249<br>(0.00016) | -1.068 | -1.429 | -1.550 |
| ***COX1*** |  |  |  |  |  |  |  |  |
| All | 151 | 628 | 15 | 0.614 (0.033) | 0.00517<br>(0.00023) | -0.1242 | -1.797 | -1.381 |
| CH-North | 92 | 628 | 7 | 0.167 (0.053) | 0.00031<br>(0.00011) | -2.039* | -3.117* | -3.264** |
| CH-South | 59 | 628 | 8 | 0.485 (0.078) | 0.00103<br>(0.00021) | -1.468 | -1.179 | -1.497 |
| ***concatenated cytb + COX1*** |  |  |  |  |  |  |  |  |
| All | 141 | 1076 | 31 | 0.792 (0.029) | 0.00658<br>(0.00026) | -0.067 | -2.848 | -2.034 |
| CH-North | 86 | 1076 | 17 | 0.557 (0.065) | 0.00080<br>(0.00014) | -2.209*** | -3.415** | -3.545** |
| CH-South | 55 | 1076 | 14 | 0.706 (0.064) | 0.00157<br>(0.00018) | -1.194 | -2.147 | -2.158 |

\* $p < 0.05$ , \*\* $p < 0.02$ , \*\*\* $p < 0.0$

To establish phylogeographic patterns, we selected a subset of the samples collected; 2-9 eggs were collected from freshly-laid clutches in 72 populations of *R. temporaria* in the Swiss Alps during the 2013 breeding season. Locations, elevations, and sample sizes of the populations are given in Table S1. The eggs were transported to the University of Zürich, where they were reared at 23-25 °C until they hatched and reached stage 36 (Gosner, 1960). One individual from each clutch (*N* = 368) was euthanised and stored in ethanol.

### Molecular lab work

Total DNA was extracted from tadpole tails by overnight digestion in 10% proteinase K solution at 56 °C and extracted using the Qiagen Biosprint 96 DNA blood kit (Qiagen, CA, USA). DNA was eluted in 100 - 200 μL buffer AE (QIAgen).

#### mitochondrial DNA

To establish phylogeographic patterns at a European scale, we sequenced a 448bp fragment of *cytb* to correspond with previous studies (Teacher *et al.* 2009; Vences *et al.* 2013). The gene was amplified by PCR using *Rana-cytb-F2* (5’ TTAGTAATAGCCACAGCTTTTGTAGGC), and *Rana-Cytb-R2* (5’ AGGGAACGAAGTTTGGAGGTGTGG) primers (Vences *et al.* 2013) with an annealing temperature of 53 °C.

The aforementioned studies do not include fine-scale sampling of the Italian Alps. To determine the phylogenetic relationship between Swiss and Italian samples generated sequence data to compare with Italian samples from Stefani *et al.* (2012). For a subset of 44 populations (151 families), we PCR-amplified a 628bp fragment of COXI. Primers L4437 (5’ AAGCTTTCGGGCCCATACC) and H6564 (5’ GGGTCTCCTCCTCCAGCTGGGTC) with an annealing temperature of 49 °C.

Amplified COXI and *cytb* fragments were subjected to Sanger sequencing after we purified them using a standard ExoSap protocol: 0.25 μl Exonuclease I (20 U/μl; New England Biosystems), 0.5 μl rAPID Alkaline Phosphatase (SAP) (1 U/μl), 7.25 μl nuclease free water, 8 μl PCR product. Cycling conditions were as follows: 37 °C for 45 min, 80 °C for 15 min, and held at 10 °C. From the cleaned product, 2 μl per sample was used in 10 μl sequencing reactions (BigDye Sequencing Kit, Applied Biosystems). Samples were sequenced in one direction for each gene using an automated 3130xl DNA Analyzer (Applied Biosystems, CA, USA) and the sequences were aligned using CLC Main Workbench 5.0.2 (CLC Bio) and BioEdit (Hall 1999). Sequences were verified as *cytb* and COXI using the BLASTN 2.2.24 algorithm (Altschup *et al.* 1990) implemented on the National Center for Biotechnology Information (NCBI) online platform.

Palo *et al.* (2004) identified a restriction enzyme cut site at position 277 in the *cytb* sequence that distinguishes between the eastern and the western mitochondrial clades. We used this assay to assign our samples to clades. Restriction digests were performed using 8 μl of the PCR product, 0.1 μl 10000 U/ml *StyI* restriction enzyme (New England Biolabs), 0.9 μl water, and 1 μl 10X Buffer (Qiagen). PCR products were digested for 30 minutes at 37 °C, where after they were resolved on a 1% agarose gel and assessed by eye under UV-light.

### Analysis

#### Geographic distribution of mitochondrial variation

To visualise the geographic distribution of genetic variation, haplotype networks were constructed with *cytb* and COX1 sequences using statistical parsimony (Templeton *et al.* 1992) as implemented in TCS version 1.21 (Clement *et al.* 2000). Within Switzerland, data generated in this study were used: 368 individuals sequenced at 448bp *cytb*, and 151 individuals sequenced at 1076bp (628bp COX1 and 448bp *cytb*). At a broader geographic scale, we visualised genetic variation by combining our data with those from elsewhere in Europe obtained from NCBI (accession numbers for *cytb*: KC799522.1 – KC800122.2; Vences *et al.* 2013). This dataset comprised 969 individuals sequenced at 331bp cytb. Note that these data do not include Italian samples described in Stefani *et al.* (2012), as a different *cytb* fragment was amplified in their study. A separate analysis was conducted based on COXI to determine the phylogenetic relationship between Swiss and Italian samples (see below).

#### Phylogenetic analyses

Evolutionary relationships among geographic groups in the haplotype network were determined with phylogenetic analyses on three datasets: 1) *cytb1*, 2) COXI, 3) combined *cytb1* and COXI. All datasets comprised of data from this study and all available data from Vences *et al.* (2013). Outgroup species were chosen based on the *Rana* phylogeny in Veith *et al.* (2003) and data available from NCBI. For *cytb*, the outgroups were *R. pyrenaica* (KC799521.1), *R. arvalis* (KC800123), *R. iberica* (KC799485), and *R. italica* (KC799493). For COXI, the outgroups were *R. pyrenaica* (KC977251.1) and *R. arvalis* (JN971596.1). As a first step we identified the evolutionary model that best fit each dataset, and secondly we determine the most likely phylogenetic tree under the chosen model. Phylogenetic trees were estimated using both Bayesian and likelihood methods.

The most likely evolutionary mutation model was determined for each dataset by comparing 88 alternative models in jModeltest2 (Darriba *et al.* 2012) and using the Akaike Information Criterion (Akaike 1973) to select the model best supported by the data.

Phylogenetic analyses were conducted using maximum likelihood and Bayesian methods. Maximum likelihood analyses were implemented in RAxML v.8.2.2 (Stamatakis 2006). For each dataset, 1000 rapid bootstrap replicates were conducted followed by a maximum likelihood search for the most likely phylogenetic tree. Bayesian phylogenetic analyses were conducted using MrBayes v.3.2.1 (Huelsenbeck & Ronquist 2001). Analyses were run for 1 × 10^6^ generations and sampled every 1000 generations. Four Metropolis-coupled Markov chain Monte Carlo (MCMCMC) chains were used for two replicate runs of each dataset, with chain heating kept at default temperatures. Convergence was assessed using the diagnostic output from the *sump* command: Adequate effective samples sizes (ESS) for each parameter (>200), appropriate mixing of chains, and an average standard deviation of split frequencies between independent runs <0.05. A burnin of 25% was used to obtain the consensus phylogram and posterior probabilities for each bipartition.

Nomenclature for the major lineages determined either side of the Alpine ridge will be CH-North (northern Swiss lineage) and CH-South (southern Swiss lineage). A third lineage referred to as CH-brown is phylogenetically closer to the CH-North lineage, but occurs exclusively south of the bisecting alpine ridge. To estimate the geographic distribution of genetic diversity, haplotype diversity (H) and nucleotide diversity (π) were calculated in DnaSP 5.0 (Librado & Rozas 2009) for all Swiss populations and for CH-North and CH-South separately.

We estimated the coalescence date for all *R. temporaria* haplotypes and the Swiss subset of haplotypes using a Bayesian Markov chain Monte Carlo analysis with *R. arvalis* and *R. pyrenaica* as outgroups (BEAST v. 1.8.2; Drummond *et al.* 2012). COX1 and *cytb1* were analysed as a single dataset, totaling 1076bp, because jModeltest inferred a similar nucleotide substitution model for both gene fragments. We used the GTR + I substitution model, with estimated base frequencies, and 6 rate categories. Priors were set to their defaults except for the date of divergence of the three *Rana* species, which was specified as a lognormal distribution with a mean of 3.18 Mya and standard deviation of 0.43 Mya (Veith *et al.* 2003). Tests were run with all demographic and molecular clock combinations as priors, and the most likely model was selected based on a marginal likelihood test. The final analysis was run using a relaxed molecular clock, and a demographic model of constant population size.

To test the hypothesis that Switzerland was colonised from the Italian refugium, we estimated the phylogenetic relationship between 569 bp COX1 data from this study, and data available on NCBI. This analysis was based on COX1 data only, because the publicly available *cytb* fragment sequenced for samples from northern Italy (Stefani *et al.* 2012) does not overlap with that of samples from Europe (Vences *et al.* 2013). We included *R. temporaria* haplotypes from the Italian Alps (Stefani *et al.* 2012; FN813783-FN813812), Europe (Vences *et al.* 2013; KC977228.1 - KC977251.1) and outgroups *R. pyrenaica* (KC977251.1) and *R. arvalis* (JN971596.1). Phylogenetic analyses were conducted as described above.

#### Demographic history

To test for signals recent changes in population size we used Tajima’s D, Fu’s F, and the DNA sequence mismatch distribution (Rogers & Harpending 1992). A mismatch distribution tests for signatures of population expansion based on the number of nucleotide differences (mismatches) between DNA sequences. When population size remains stable the distribution of mismatches is expected to be multimodal, reflecting the stochastic nature of gene trees. When populations expand, we expect a unimodal distribution of mismatches. Signals of population expansion were tested in the haplogroups using Harpending’s raggedness index (RI). The expansion time was estimated under a model of pure demographic expansion with parameters set to default in Arlequin (Excoffier & Lischer 2010). The parameter of demographic expansion, Tau, was estimated according to Schneider & Excoffier (1999). The validity of the expansion model was tested using the sum of square deviations (SSDs) between the observed and expected mismatches as implemented in Arlequin.

## Results

The final dataset comprised 368 families from 72 populations sequenced at 448bp of the *cytb* gene, and 151 families from 44 populations sequenced at 628bp of COXI. Based on the absence of the diagnostic *cytb* restriction site, all Swiss samples were determined to be from the western clade.

### Geographic distribution of genetic variation

#### Haplotype network

Overall the haplotype network shows an eastern (blue) and western (other colours) clade with much higher genetic diversity found in the western clade (Fig. 1). The Swiss samples assigned to three western haplogroups; a green haplogroup which is widespread across western Europe and Great Britain, a brown haplogroup found almost exclusively in the Pyrenees (Vences et al. 2013), and a Swiss specific haplogroup shown in purple described here for the first time. Within Switzerland the green haplogroup is found north of the Alpine ridge bisecting Switzerland, while the purple and brown haplogroups occur south of the ridge (Fig. 2). This division was confirmed when Swiss samples were analysed using cytb alone and in combination with COXI (Fig. S2.1). The haplogroup found north of the Alps (CH-North) corresponded with the widespread western European haplogroup, which is thought to have colonised western Europe from the Pyrenees (green haplotypes in Fig. 1; Vences *et al.* 2013). The brown haplogroup (CH-brown) was closely related to the common western European haplogroup, and occurs almost exclusively in the Pyrenees (brown haplotypes in Fig. 1). The southern haplogroup was unique to Switzerland, and fell between the haplogroups sampled in the Pyrenees and those sampled in Croatia and Greece in the haplotype network. CH-North and CH-South co-occurred in some populations but across all elevations in eastern Switzerland and at low elevation in western Switzerland. A second western European lineage (CH-brown: brown haplotypes in Fig. 1), found in the Pyrenees and restricted roughly to the southern part of Europe, co-occurred with CH-South haplotypes in southern Switzerland (Fig. 2).

**Figure 2.**
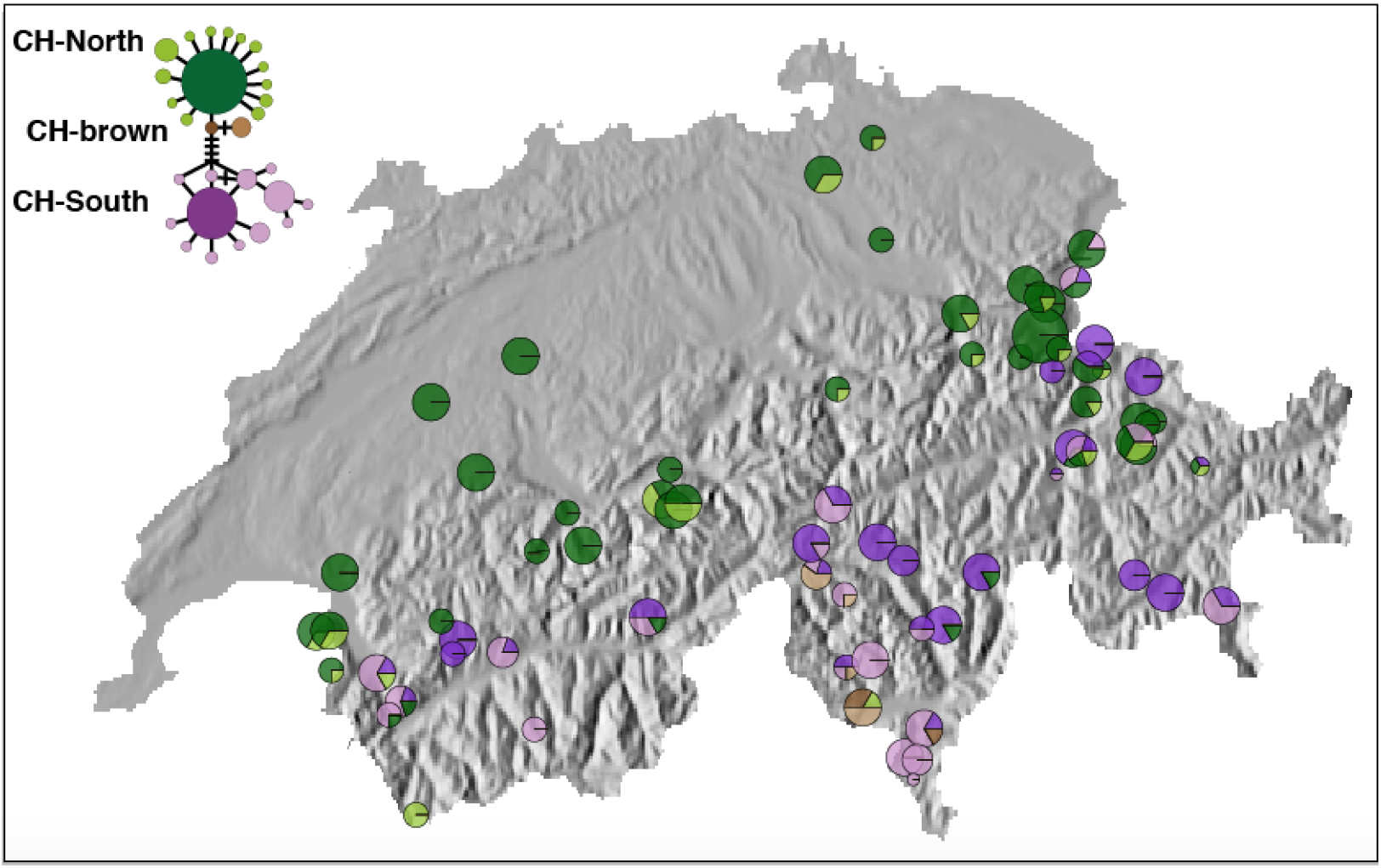
Swiss *cytb* (448bp) haplotypes are distributed roughly north (CH-North) and south (CH-South and CH-brown) of the northern Alps. Pie charts show the proportion of individuals that had northern or southern haplotypes in each population, with pie chart size reflecting the number of samples per population. The inset depicts the *cytb* haplotype network calculated for all Swiss samples (368 individuals from 72 populations sequenced at 448 bp). The most common haplotype within each haplogroup is shaded in a darker colour compared to the derived haplotypes. Thus the distribution of the common and derived haplotypes are shown with pie charts on the map.

#### Phylogenetic analyses

The same phylogenetic relationships were recovered using ML and Bayesian approaches, as well as between the *cytb1*, COXI, and concatenated datasets, and were concordant with previously published phylogenies (Fig. 3) (Veith *et al.* 2002; Vences *et al.* 2013). A basal split in the *R. temporaria* clade separates three Spanish clades that occur exclusively in Iberia (Vences *et al.* 2013) from the rest of the samples. *R. temporaria* is further split into an eastern and a western clade with high posterior probability (0.99). Within the western clade, a further split is found between populations from southern Switzerland and all other western populations (including northern Switzerland/western Europe; posterior probability = 0.97).

**Figure 3.**
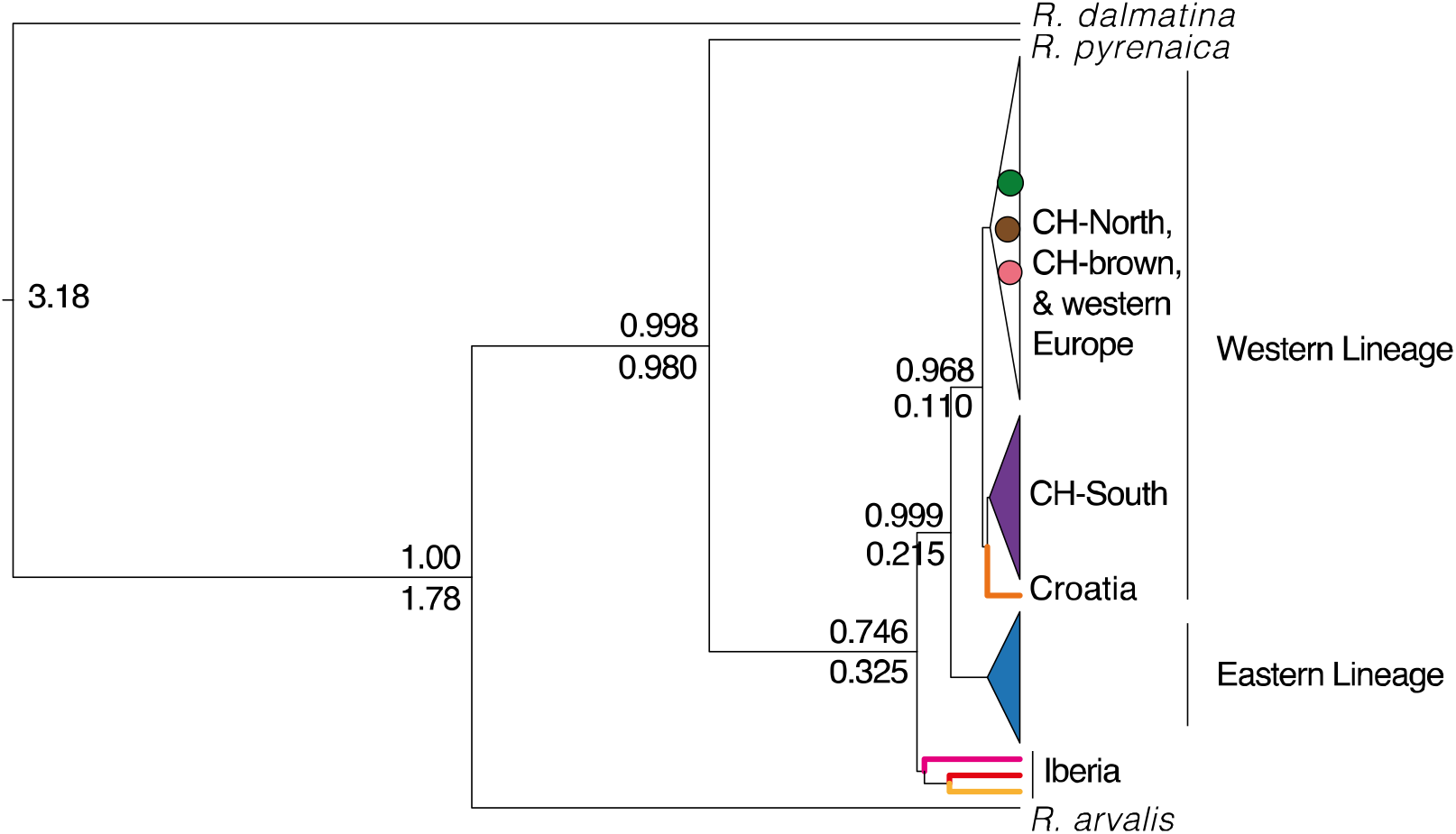
Bayesian phylogenetic tree of concatenated *cytb1* and COXI genes. The eastern and western lineage, the CH-South, CH-North, and CH-brown haplogroups are defined as in Fig. 1. The eastern and CH-South haplogroups are coloured as in Fig. 1. CH-North, CH-brown and western European haplotypes are indicated by green, brown, and pink circles (colours as in Fig. 1). Posterior probability is shown above the line, and the inferred coalescent time (Mya) as determined by BEAST is shown below the line. The calibration used (3.18Mya; Vences *et al.* 2003) is shown at the root of the tree.

The estimated mean coalescent times in Fig. 3 support a divergence between the eastern and western clades at ~210 kya, and a more recent divergence between CH-South and the rest of the western clade at ~110 kya. The most likely coalescent model based on marginal likelihood tests was a relaxed molecular clock and constant population size. The effective sample size (ESS) for all parameters was above 200, and all independent runs showed convergence and stationarity.

#### Distribution of genetic variation

Genetic diversity within Switzerland was higher in populations found south of the alpine ridge bisecting Switzerland. Haplotype and nucleotide diversity based on 448bp *cytb* gene fragment was roughly double in CH-South when compared to CH-North populations, and roughly three times higher based on 628bp COX1 sequence (Table 1; diversity measures for individual populations are reported in Tables S1 and S2).

### Demographic history

The mismatch distribution analyses did not converge for CH-North for any of the datasets, but Tajima’s D and Fu’s F were negative and significant for all CH-North datasets indicative of a population expansion. Tajima’s D and Fu’s F were non-significant for CH-South, and there was no evidence for population expansion for any of the CH-South analyses.

### Origin of haplotypes in Switzerland

Phylogenetic analysis based on 569 bp of COX1 revealed that haplotypes from the Italian Alpine Lineage (AP) I described in Stefani *et al.* (2012) grouped with CH-South haplotypes with high posterior probability (0.94; Fig. 4). Three haplotypes identified by Stefani *et al.* (2012) – FN813810.1, FN813785.1, FN813786 – were identical to CH-South haplotypes. API occurs in the central Italian Alps and contains the most abundant Italian Alpine haplotype (CA2), which was identical to CHS03. In our sample, this haplotype was found only in a population on a high pass in southern Switzerland, on the border with Italy. The most widespread CH-South haplotype was identical to the widespread API haplotype DE10. A third haplotype (API: VF6) was identical to CHS06, found only at high elevation in southwestern Switzerland (Figs. 2.4 & 2.5).

**Figure 4.**
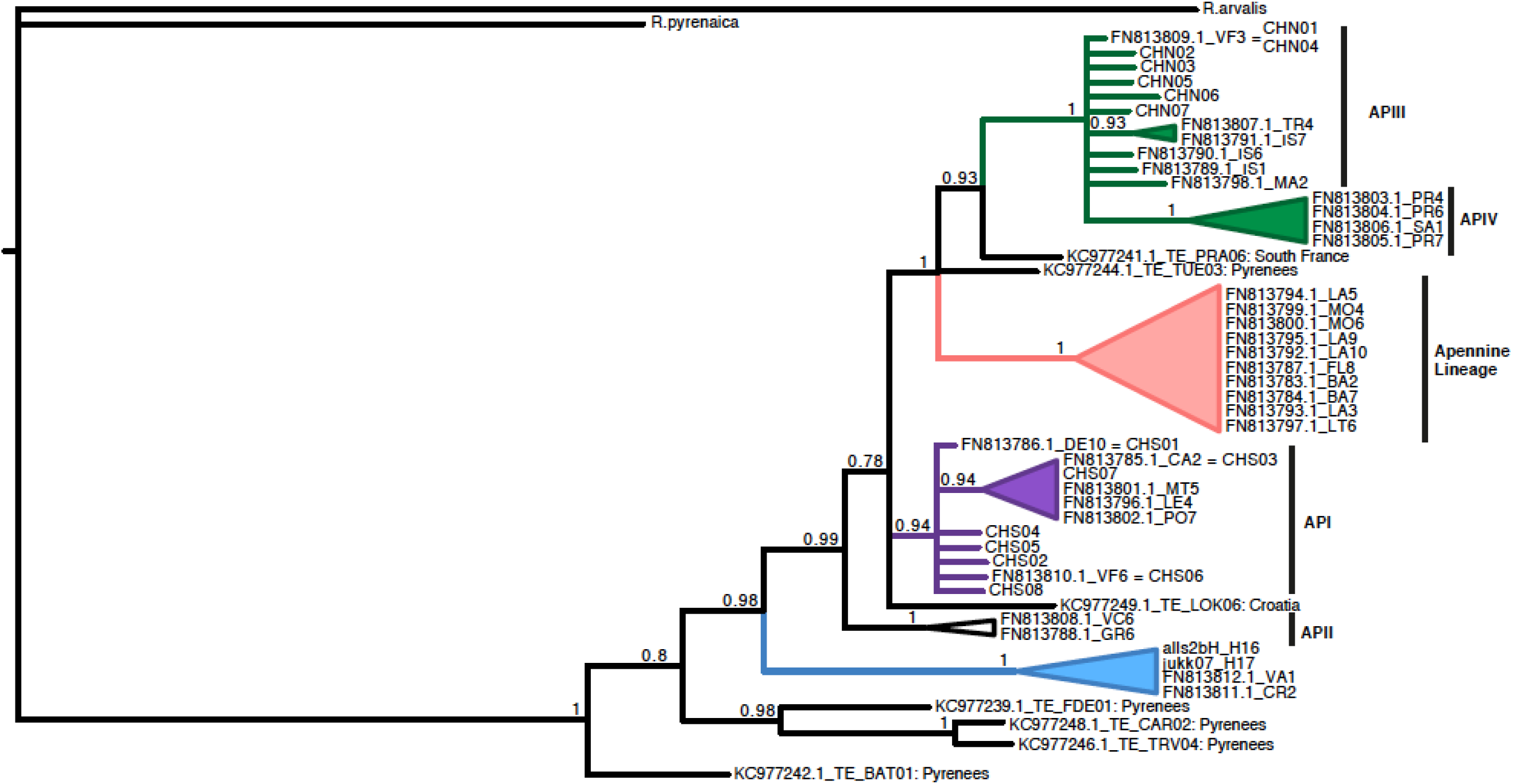
Bayesian phylogenetic tree of COX1 haplotypes (569 bp) sampled across the *R. temporaria* range. Italian lineages described by Stefani *et al.* (2012) (Alpine lineage I-IV and Apennine lineage; NCBI code FNxx) are shown with vertical black bars. Four haplotypes from this study match Stefani *et al.* (2012) haplotypes and are indicated with an equal (=) sign. Coloured regions match the haplogroups identntif ied in Fig.1. Haplotypes from this study are indicated with CHN (CH-North) and CHS (CH-South). CH-brown haplotypes are not visible in this figure as these populations were not sequenced at COX1. Haplotypes from Vences *et al.* (2013) are shown with their NCBI codes (KCxx) and their sample locations. Posterior probability of >0.5 is shown above each node.

The northern Swiss lineage (CH-North) clustered with APIII with high posterior probability (1.0). One previously identified haplotype from APIII (FN813809.1 = VF3) was identical to the most common CH-North haplotype (CHN01). Geographically, APIII and VF3 are found in northwestern Italy, but in Switzerland CHN01 occurs mostly north of the bisecting alpine ridge (Fig. 5). APIII clusters with a haplotype from southern France with high posterior probability (0.93; Fig. 4), which suggests that this haplogroup is analgous to the green haplogroup described by Vences *et al.* (2013) and is widespread across France, Germany, and Great Britain.

**Figure 5.**
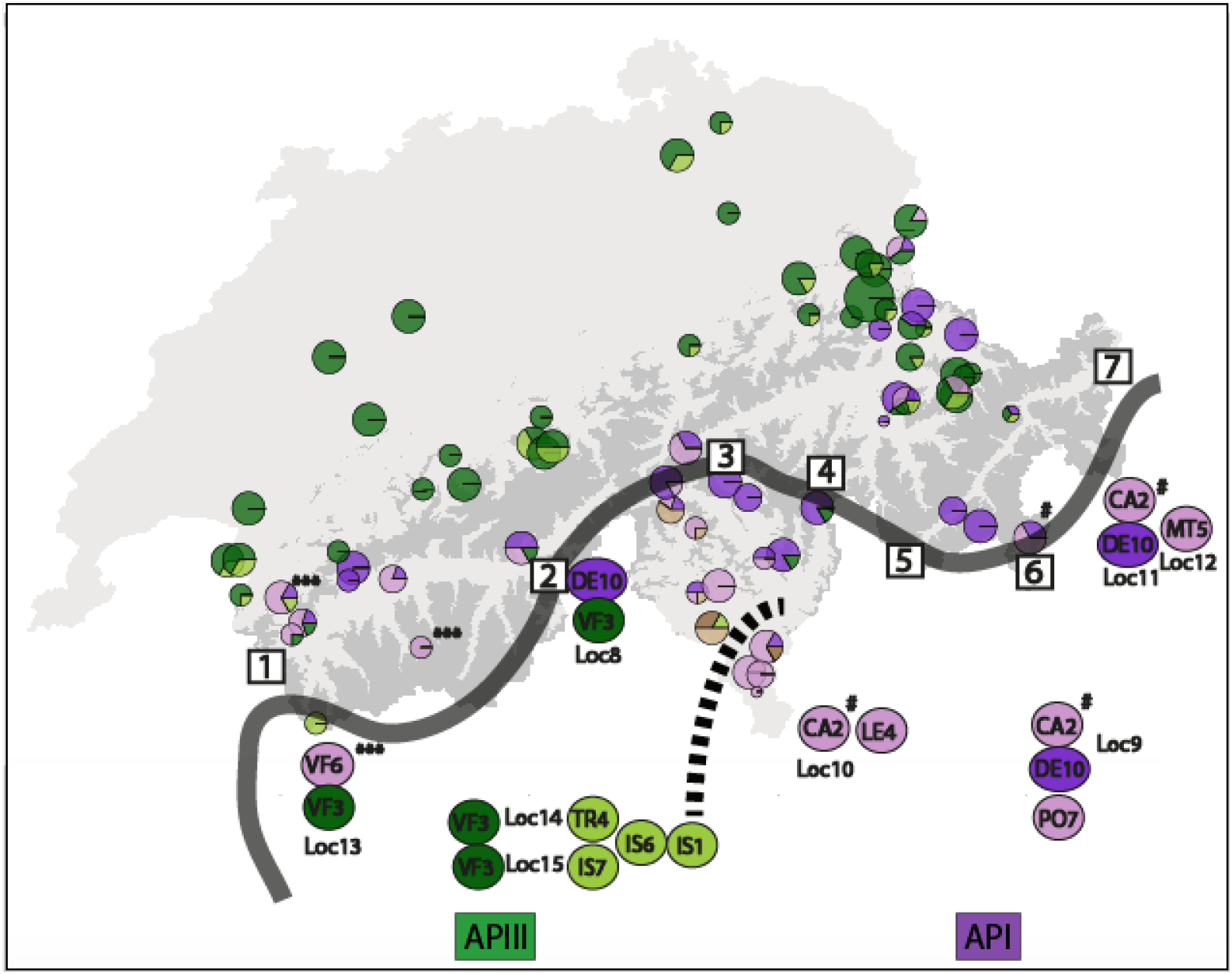
Possible trans-alpine colonisation routes from the Italian Alps into Switzerland. The map shows elevation >2000 m in dark grey; thus the light grey highlights the potential regions available for colonising populations. The thick solid grey line roughly denotes the central Alpine ridge. Haplotype colours correspond with the haplogroups identified in Fig. 2 and 4. We show the geographic distribution of the COX1 haplotypes identified by Stefani *et al.* (2012) in Northern Italy bordering Switzerland. The dashed line indicates the Ticino valley that roughly separates API (light and dark purple) from APIII (light and dark green). For illustration purposes we show the *cytb* haplotypes from Switzerland (i.e. Fig. 3) as mitochondrial haplotypes are linked and this gene was sequenced more geographically extensively than COX1 for our study. The Italian populations are defined as in Stefani *et al* (2012); Loc8: Pramollo (46.36 N 11.86 E), Loc9: M.Pora (45.88 N 10.1 E), Loc10: Lemna (45.85 N 9.18 E), Loc11: S.Caterina (46.41 N 10.49 E), Loc12: V.Martello (46.49 N 10.69 E), Loc13: V.Ferret (45.85 N 7.02 E), Loc14: Traversella (45.53 N 7.72 E), Loc15: Issiglio (45.46 N 7.32 E). Four haplotypes matched sequences from this study and their geographic locations are shown as follows: VF3: Dark green; DE10: Dark purple; CA2:#; VF6:***. The numbered blocks are mountain passes less than 2000 m elevation presenting possible cross-alpine colonisation points: 1=Col de Forclaz, 2=Simplonpass, 3=Lukmanierpass, 4=San Bernardino Pass, 5=Malojapass, 6=Bernina Pass, 7=Reschenpass.

## Discussion

We interpret our results to indicate that *Rana temporaria* recolonised western Europe almost exclusively from multiple refugia in the Italian Alps. Populations originating from hypothesised refugia in the Iberian Peninsula occur only in Spain and southwestern France. Fine-scale sampling of populations across Switzerland provides evidence of a trans-alpine colonisation from the central Italian Alps into Switzerland, and recolonisation from the western Italian Alps around the alpine arc into much of western Europe. We found two deeply diverged mitochondrial lineages occurring roughly north and south of the northern Alps, with contact zones in eastern and western Switzerland. These results strongly suggest that that the Alps did not present a significant barrier to dispersal for this species during the recolonisation of Europe. This has resulted in complex regional variation in the genetic structure of *R. temporaria* across Switzerland, which could have important implications for population persistence and adaptive potential.

## Eastern and Western European lineages

Consistent with previous research, we found a deep phylogenetic split between eastern and western European lineages of *R. temporaria* (Palo *et al.* 2004). The age of the east-west divergence, calibrated based on the *R. dalmatina* divergence time (3.18 Mya; Veith *et al.* 2003), was placed in the late Pleistocene (~0.215 Mya). The divergence date was previously estimated as ~0.71 Mya (CI 0.53-0.95 Mya) based on *cytb* sequences (Palo *et al.* 2004), corresponding to the onset of strong Pleistocene climatic cycles. The discrepancy may be attributed to the large confidence intervals associated with the statistical priors, as well as the slightly different calibration points included in both studies. However, our estimated date and that of Palo et al.(2004) both suggest that the divergence between the eastern and western clades is attributable to the dramatic climatic cycles that occurred during the Pleistocene, which began ~1.8Mya.

Two previous studies disagreed about the existence of a contact zone between the eastern and western lineages in northern Switzerland and along the French Mediterranean into Spain (Teacher *et al.* 2009, Rodrigues et al. 2013). Our thorough sampling of northern Switzerland, including several populations in the lowlands where the contact zone was said to lie (Teacher et al. 2009), uncovered no eastern haplotypes whatsoever and hence no evidence of contact between clades. Presumably, a secondary contact zone between the eastern and western haplogroups occurs somewhere farther to the east.

## Post-glacial recolonisation of western Europe

Several studies have identified multiple glacial refugia in the Italian peninsula, that consequently lead to multiple deeply diverged lineages (Canestrelli *et al.* 2006, 2008; Crottini *et al.* 2007; Canestrelli & Nascetti 2008). Our results support the assertion that *R. temporaria* re-colonised western Europe from multiple sub-refugia within the Italian refugium (Stefani *et al.* 2012). We identified two western mitochondrial lineages distributed roughly north and south of the northern Alps within Switzerland. The two lineages are estimated to have diverged ~110 kya, at the onset of the most recent glacial cold period. Four Alpine lineages have been described in northern Italy, two of which (API and APIII) are distributed south of Switzerland roughly east and west of the Ticino valley (Fig. 5). Our phylogenetic analyses suggest that the southern (CH-South) haplogroup was most likely colonised by API from the central Italian Alps, while the northern (CH-North) haplogroup was colonised by APIII.

A thorough geographic analysis of the CH-North and CH-South lineages suggests that they recolonised western Europe via two different routes (Fig. 6). The geographic distribution of the southern lineage across the central Alpine ridge is evidence of multiple trans-alpine colonisations into Switzerland (Fig. 5). Three CH-South haplotypes were identical to AP1 haplotypes, and their geographic distribution can be used to narrow down colonisation routes. Assuming a conservative maximum habitable elevation of 2000 m (as in Braaker & Heckel 2009), there are seven possible passes across the central Alpine ridge (Fig. 5). The most widespread AP1 haplotype (CA2) was identical to a CH-South haplotype found in south-eastern Switzerland, which suggests colonisation across the Maloja Pass (5, Fig. 5) or Bernina Pass (6, Fig. 5).

**Figure 6.**
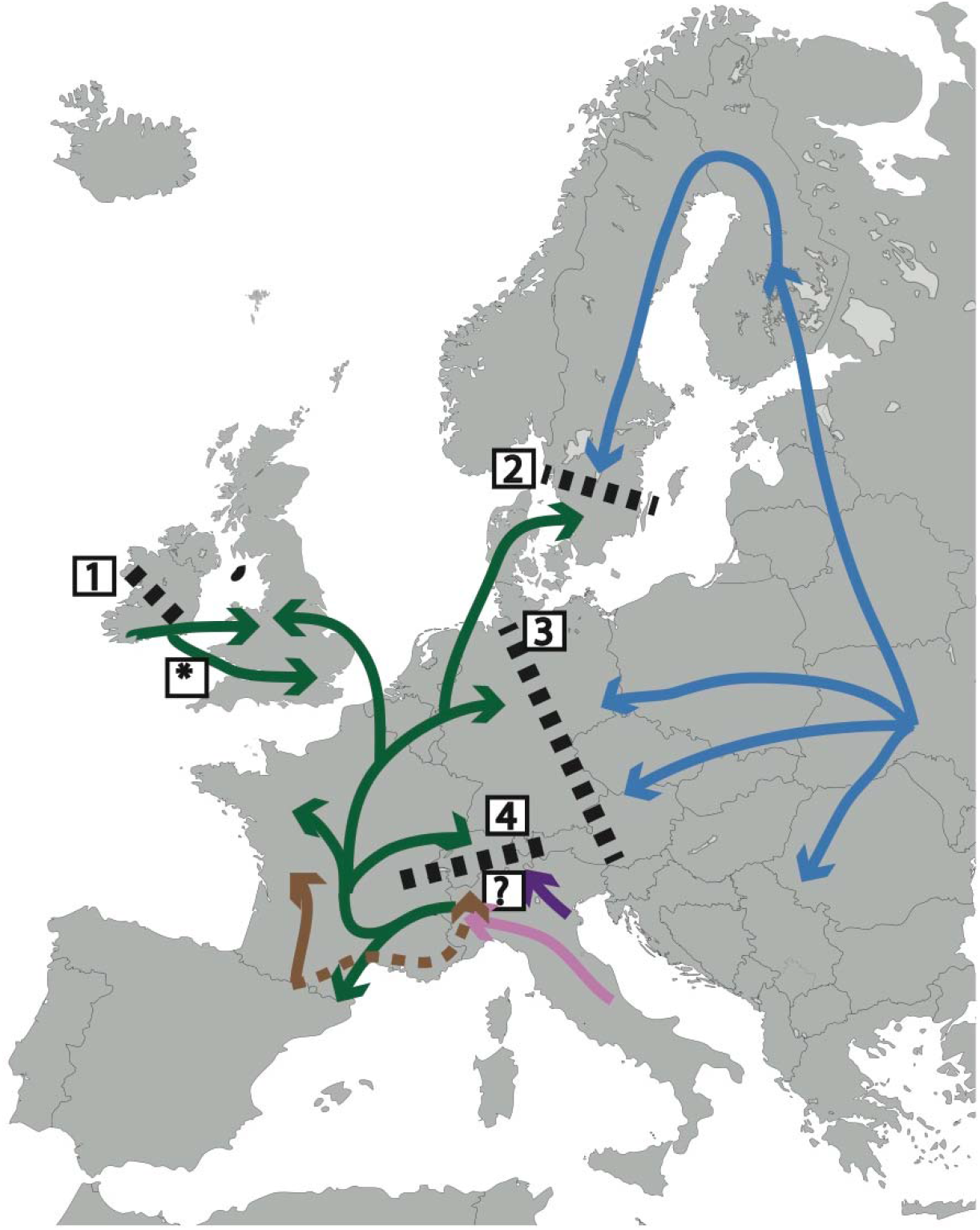
Proposed *R. temporaria* post-glacial recolonisation routes. Coloured arrows indicate the proposed origin and colonisation routes of five haplogroups. Our results suggest that Italy is the major glacial refugia for *R. temporaria*, with the common green haplogroup colonising western Europe from the north western Italian Alps, and a transalpine colonisation of the purple haplogroup into Switzerland. Dashed lines show probable regions of secondary contact between diverged lineages. 1=common western European lineage meets *R. temporaria* from Irish refugium (Teacher *et al.* 2009); 2=Likely original contact zone between eastern and western haplogroup, but only the western haplogroup is currently found here (Palo *et al.* 2004); 3=Conact zone between eastern and western haplogroup in northern Germany (Schmeller *et al.* 2008), but contact zone south of this site still needs to be described; 4=contact zone between CH-South and CH-North described in this paper; ?=Possible contact zone between the Iberian (CH-brown) and Italian lineages described in this paper. The asterist (*) denotes an alternative origin and colonisation route of the green haplotype from the Irish refugium into western Europe.

Southwestern Switzerland contained several sites with secondary contact between API and APIII, which could suggest a southern trans-alpine colonisation route of APIII into Switzerland. Possible routes include the Col de Ferdaz (1, Fig. 5) in the west or Simplonpass (2, Fig. 5), but the most likely point of colonization is difficult to identify because the most common CH-South haplotype (API haplotype DE10) was geographically widespread in northern Italy.

In other European taxa, studies of trans-alpine colonisation have identified the same trans-Alpine colonisation routes for both plants and animals (Lugon-Moulin & Hausser 2002; Yannic *et al.* 2008, 2012; Braaker & Heckel 2009; Parisod 2008). The example of European white oaks (Mátyás & Sperisen 2001), which presently occur only rarely above 1200 m in the Alps (Rigling *et al.* 2013), illustrates that trans-Alpine recolonisation was not limited only to taxa adapted to cold climates. Presumably, dispersal over Alpine passes was more extensive when the treeline was ~200 m higher than it is today, which occurred regularly during the period of 5’000-10’000 years ago (e.g. Ammann *et al.* 2000; Tinner & Theurillat 2003; Heiri *et al.* 2006). Exact colonisation routes could be investigated with the inclusion of nuclear markers and fine-scale geographic sampling of populations across the putative passes used for colonisation.

CH-North haplotypes appear to have colonized Switzerland from the west probably up the Rhone River or from the north west over the Jura Mountains (Fig. 6). The geographic distribution of the derived CH-North haplotypes towards southern Switzerland (light green haplotypes; Fig. 2) and overall low genetic diversity (Table 1) suggest that this lineage expanded southwards in Switzerland rather than colonising via trans-alpine routes as with the CH-South lineage. Three possible origins of this haplogroup have been proposed: Iberia, a cyptic refugium in southern Ireland, and northern Italy (Teacher *et al.* 2009; Stefani *et al.* 2012; Vences *et al.* 2013). Although we cannot definitively rule out Iberia or Ireland, our results seem to suggest an Italian origin. Specifically, the previously identified common western European haplogroup (T4; Vences *et al.* 2013) occupies northern Switzerland, and we found that this haplogroup is identical to Italian APIII found in the north western Italian Alps. In addition, the rest of the CH-North haplotypes form a monophyletic group with APIII (Fig. 4). Finally, genetic diversity is expected to decrease with distance from glacial refugia as populations experience bottleneck events with colonisation (Hewitt 1996). The genetic variation in the green/APII/T4 haplogroup is substantially higher in northern Italy than in Ireland or Iberia, suggesting that this was the main refugium of the haplogroup.

The identification of a second CH-brown lineage exclusively south of the central Alpine ridge in southern Switzerland suggests that there may be eastward expansion from Iberia south of the Alps (Fig. 2 & 2.5). This lineage is ubiquitous in the Pyrenees, and is found in southern France, northern Italy, and Croatia (Vences *et al.* 2013). Deglaciation of the Tinee Valley on the border between France and Italy occurred during the Oldest Dryas (~15kya), earlier than the rest of the Alps where glaciers retreated during the Younger Dryas (~11kya) and the late ice retreat (~9kya) (Darnault *et al.* 2011). This potential early colonisation corridor could have provided the way for an eastward expansion from the Pyrenees, across northern Italy, to Croatia. Although westward expansion from the Carpathian refugium through northern Italy has been described (Demesure *et al.* 1996; Taberlet *et al.* 1998), we found no examples of an eastward expansion from Iberia into northern Italy. Longer mtDNA fragments and more extensive sampling would be needed to investigate this unusual colonisation route further.

## Conclusions

The *R. temporaria* western mitochondrial clade is exceptional in that it harbours more genetic diversity than any other European brown frogs (Palo *et al.* 2004; Vences *et al.* 2013). We show that much of this variation can be attributed to the European Alps. The topographic complexity of the Alps has resulted in multiple glacial refugia and subsequent divergence between *R. temporaria* lineages. After the retreat of the glaciers, the Alps presented a semi-permeable barrier to dispersal for these lineages, resulting in multiple colonisation events in the south. However, this also limited colonisation from other refugia in northern Italy, which subsequently followed a circum-Alpine route to colonise western Europe and Switzerland (Fig. 6). This has resulted in two deeply diverged lineages that occur in the north and the south of Switzerland with contact zones in the east and west. Our work highlights how fine-scale phylogeographic studies across the Alps can elucidate of the phylogeographic history of cold-adapted species in Europe.

## Author contributions

AJvR and JVB conceived of the project. AJvR, MR, and SB generated and analysed the data. All authors contributed to the writing of the manuscript. This manuscript forms part of AJvR’s PhD thesis.

## Data accessibility

All sequences are available on NCBI. Accession numbers for *cytb*: MF624310-MF624355; COX1:

## References

Akaike H (1973) Information theory and an extension of the maximum likelihood principle. In: 2nd International Symposium on Information Theory, Tsahkadsor, Armenia, USSR, pp. 267–281. Budapest:Akadémiai Kiadó.

Alberto F, Niort J, Derory J et al. (2010) Population differentiation of sessile oak at the altitudinal front of migration in the French Pyrenees. Molecular Ecology, 19, 2626–2639.

Alton LA, Franklin CE (2017) Drivers of amphibian declines: effects of ultraviolet radiation and interactions with other environmental factors. Climate Change Responses, 4, DOI 10.1186/s40665-017-0034-7.

Altschup SF, Gish W, Miller W, Myers EW, Lipman DJ (1990) Basic Local Alignment Search Tool. Journal of Molecular Biology, 205, 403–410.

Alvarez N, Thiel-Egenter C, Tribsch A et al. (2009) History or ecology? Substrate type as a major driver of spatial genetic structure in Alpine plants. Ecology Letters, 12, 632–640.

Ammann B, Birks HJB, Brooks SJ et al. (2000) Quantification of biotic responses to rapid climatic changes around the Younger Dryas — a synthesis. Paleogeography, Paleoclimatology, Paleoecology, 159, 313–347.

Avise JC, Riddle B (2009) Phylogeography◻: Retrospect and Prospect. Journal of Biogeography, 36, 3–15.

Bachmann JC (2017) Adaptive Divergence across an Elevational Gradient in the Common Frog (Rana temporaria). University of Zurich.

Bolger AM, Lohse M, Usadel B (2014) Trimmomatic: A flexible trimmer for Illumina sequence data. Bioinformatics, 30, 2114–2120.

Bonin A (2008) Population genomics: a new generation of genome scans to bridge the gap with functional genomics. Molecular ecology, 17, 3583–4.

Bonin A, Taberlet P, Miaud C, Pompanon F (2006) Explorative genome scan to detect candidate loci for adaptation along a gradient of altitude in the common frog (Rana temporaria). Molecular biology and evolution, 23, 773–83.

Borcard D, Legendre P (2002) All-scale spatial analysis of ecological data by means of principal coordinates of neighbour matrices. Ecological Modelling, 153, 51–68.

Braaker S, Heckel G (2009) Transalpine colonisation and partial phylogeographic erosion by dispersal in the common vole (Microtus arvalis). Molecular Ecology, 18, 2528–2531.

Brady LD, Griffiths RA (2000) Developmental responses to pond desiccation in tadpoles of the British anuran amphibians (Bufo bufo, B. calamita and Rana temporaria). Journal of Zoology, London, 252, 61–69.

Van Buskirk J (2012) Permeability of the landscape matrix between amphibian breeding sites. Ecology and evolution, 2, 3160–7.

Canestrelli D, Cimmaruta R, Costantini V, Nascetti G (2006) Genetic diversity and phylogeography of the Apennine yellow-bellied toad Bombina pachypus, with implications for conservation. Molecular Ecology, 15, 3741–3754.

Canestrelli D, Cimmaruta R, Nascetti G (2008) Population genetic structure and diversity of the Apennine endemic stream frog, Rana italica – insights on the Pleistocene evolutionary history of the Italian peninsular biota. Molecular Ecology, 17, 3856–3872.

Canestrelli D, Nascetti G (2008) Phylogeography of the pool frog Rana (Pelophylax) lessonae in the Italian peninsula and Sicily: Multiple refugia, glacial expansions and nuclear-mitochondrial discordance. Journal of Biogeography, 35, 1923–1936.

Cano JM, Laurila A, Palo J, Merilä J (2004) Population differentiation in G matrix structure due to natural selection in Rana temporaria. Evolution, 58, 2013–2020.

Catchen JM, Amores A, Hohenlohe P, Cresko W, Postlethwait JH (2011) Stacks: Building and Genotyping Loci De Novo From Short-Read Sequences. G3:Genes, Genomes, Genetics, 1, 171–182.

Caye K, Deist TM, Martins H, Michel O, François O (2016) TESS3: Fast inference of spatial population structure and genome scans for selection. Molecular Ecology Resources, 16, 540–548.

CH2014–Impacts (2014) Toward quantitative scenarios of climate change Impacts in Switzerland. OCCR, FOEN, MeteoSwiss, C2SM, Agroscope, and ProClim, Bern, Switzerland.

Clement M, Posada D, Crandall KA (2000) TCS◻: a computer program to estimate gene genealogies. Molecular Ecology, 9, 1657–1659.

Cornetti L, Lemoine M, Hilfiker D et al. (2016) Higher genetic diversity on mountain tops: the role of historical and contemporary processes in shaping genetic variation in the bank vole. Biological Journal of the Linnean Society, 118, 233–244.

Croteau MC, Davidson MA, Lean DRS, Trudeau VL (2008) Global Increases in Ultraviolet B Radiation◻: Potential Impacts on Amphibian Development and Metamorphosis. Physiological and Biochemical Zoology, 81, 743–761.

Crottini A, Andreone F, Kosuch J et al. (2007) Fossorial but widespread: the phylogeography of the common spadefoot toad (Pelobates fuscus), and the role of the Po Valley as a major source of genetic variability. Molecular Ecology, 16, 2734–2754.

Danecek P, Auton A, Abecasis G et al. (2011) The variant call format and VCFtools. Bioinformatics, 27, 2156–2158.

Dansgaard W, Johnsen SJ, Clausen HB et al. (1993) Evidence for general instability of past climate from a 250-kyr ice-core record. Nature, 364, 218–220.

Darnault R, Rolland Y, Braucher R et al. (2011) Timing of the last deglaciation revealed by receding glaciers at the Alpine-scale◻: impact on mountain geomorphology. Quaternary Science Reviews, 1–16.

Darriba D, Taboada GL, Doallo R, Posada D (2012) jModelTest 2: more models, new heuristics and parallel computing. Nature Methods, 9, 772.

Debieu M, Tang C, Stich B et al. (2013) Co-Variation between seed dormancy, growth rate and flowering time changes with latitude in Arabidopsis thaliana. PloS one, 8, 1–12.

Demesure B, Comps B, Petit RJ (1996) Chloroplast DNA Phylogeography of the Common Beech (Fagus sylvatica L.) in Europe. Evolution, 50, 2515–2520.

Drummond AJ, Suchard MA, Xie D, Rambaut A (2012) Bayesian Phylogenetics with BEAUti and the BEAST 1.7. Molecular Biology and Evolution, 29, 1969–1973.

Dudaniec RY, Spear SF, Richardson JS, Storfer A (2012) Current and historical drivers of landscape genetic structure differ in core and peripheral salamander populations. PloS one, 7.

Eaton DAR (2014) PyRAD: Assembly of de novo RADseq loci for phylogenetic analyses. Bioinformatics, 30, 1844–1849.

Ellis N, Smith SJ, Pitcher CR (2012) Gradient forests◻: calculating importance gradients on physical predictors. Ecology, 93, 156–168.

Excoffier L, Lischer HEL (2010) An Integrated Software Package for Population Genetics Data Analysis. Molecular Ecology Resources, 10, 564–567.

Fitzpatrick SW, Gerberich JC, Kronenberger JA, Angeloni LM, Funk WC (2015) Locally adapted traits maintained in the face of high gene flow. Ecology Letters, 18, 37–47.

Fitzpatrick MC, Keller SR (2015) Ecological genomics meets community-level modelling of biodiversity: mapping the genomic landscape of current and future environmental adaptation. Ecology Letters, 18, 1–16.

Forester BR, Jones MR, Joost S, Landguth EL, Lasky JR (2016) Detecting spatial genetic signatures of local adaptation in heterogeneous landscapes. Molecular Ecology, 25, 104–120.

Fox J, Weisberg S (2011) An {R} Companion to Applied Regression. Thousand Oaks CA: Sage.

Freedman AH, Thomassen HA, Buermann W, Smith TB (2010) Genomic signals of diversification along ecological gradients in a tropical lizard. Molecular ecology, 19, 3773–3788.

Frichot E, François O (2015) LEA◻: An R package for landscape and ecological association studies (B O’Meara, Ed,). Methods in Ecology and Evolution, 6, 925–929.

Frichot E, Schoville SD, Bouchard G, François O (2013) Testing for associations between loci and environmental gradients using latent factor mixed models. Molecular Biology and Evolution, 30, 1687–99.

Frichot E, Schoville S, de Villemereuil P, Gaggiotti OE, François O (2015) Detecting adaptive evolution based on association with ecological gradients: Orientation matters! Heredity, 115, 22–28.

Garcia VOS, Ivy C, Fu J (2017) Syntopic frogs reveal different patterns of interaction with the landscape: A comparative landscape genetic study of Pelophylax nigromaculatus and Fejervarya limnocharis from central China. Ecology and Evolution, 7, 9294–9306.

Gosner KL (1960) A Simplified Table for Staging Anuran Embryos and Larvae with Notes on Identification. Herpetologica, 16, 183–190.

Goudet J (2005) Hierfstat, a package for R to compute and test hierarchical F-statistics. Molecular Ecology Notes, 5, 184–186.

Gugerli F, Englisch T, Niklfeld H et al. (2008) Relationships among levels of biodiversity and the relevance of intraspecific diversity in conservation – a project synopsis. Perspectives in Plant Ecology, Evolution and Systematics, 10, 259–281.

Günther T, Coop G (2013) Robust identification of local adaptation from allele frequencies. Genetics, 195, 205–220.

Hall TA (1999) BioEdit: a user-friendly biological sequence alignment editor and analysis program for Windows 95/98/NT. Nucleic Acids Symposium Series, 41, 95–98.

Harrisson KA, Amish SJ, Pavlova A et al. (2017) Signatures of polygenic adaptation associated with climate across the range of a threatened fish species with high genetic connectivity. Molecular ecology, 26, 6253–6269.

Hecht BC, Matala AP, Hess JE, Narum SR (2015) Environmental adaptation in Chinook salmon (Oncorhynchus tshawytscha) throughout their North American range. Molecular Ecology, 24, 5573–5595.

Heiri C, Bugmann H, Tinner W, Heiri O, Lischke H (2006) A model-based reconstruction of Holocene treeline dynamics in the Central Swiss Alps. Journal of Ecology, 94, 206–216.

Hermisson J, Pennings PS (2005) Soft sweeps: molecular population genetics of adaptation from standing genetic variation. Genetics, 169, 2335–52.

Hewitt GM (1996) Some genetic consequences of ice ages, and their role in divergence and speciation. Biological Journal of the Linnean Society, 58, 247–276.

Hewitt G (1999) Post-glacial re-colonization of European biota. Biological Journal of the Linnean Society, 68, 87–112.

Hewitt G (2000) The genetic legacy of the Quaternary ice ages. Nature, 405, 907–913.

Hijmans RJ, Cameron SE, Parra JL, Jones PG, Jarvis A (2005) Very high resolution interpolated climate surfaces for global land areas. International Journal of Climatology, 25, 1965–1978.

Hitchings SP, Beebee TJC (1997) Genetic substructuring as a result of barriers to gene flow in urban Rana temporaria (common frog) populations: implications for biodiversity conservation. Heredity, 79, 117–127.

Hjernquist MB, Soderman F, Jonsson KI et al. (2012) Seasonality determines patterns of growth and age structure over a geographic gradient in an ectothermic vertebrate. Oecologia, 170, 641–649.

Hoban S, Kelley JL, Lotterhos KE et al. (2016) Finding the Genomic Basis of Local Adaptation: Pitfalls, Practical Solutions, and Future Directions. The American Naturalist, 188, 000–000.

Hoffmann A a, Sgrò CM (2011) Climate change and evolutionary adaptation. Nature, 470, 479–85.

Huelsenbeck JP, Ronquist F (2001) MRBAYES◻: Bayesian inference of phylogenetic trees. Bioinformatics Application Note, 17, 754–755.

Ilut DC, Nydam ML, Hare MP (2014) Defining loci in restriction-based reduced representation genomic data from nonmodel species: Sources of bias and diagnostics for optimal clustering. BioMed Research International, 2014, 9 pages.

Jombart T (2008) Adegenet: A R package for the multivariate analysis of genetic markers. Bioinformatics, 24, 1403–1405.

Jombart T, Ahmed I (2011) adegenet 1.3-1: New tools for the analysis of genome-wide SNP data. Bioinformatics, 27, 3070–3071.

Jones MR, Forester BR, Teufel AI et al. (2013) Integrating landscape genomics and spatially explicit approaches to detect loci under selection in clinal populations. Evolution, 67, 3455–68.

Kawecki TJ, Ebert D (2004) Conceptual issues in local adaptation. Ecology Letters, 7, 1225–1241.

Lasky JR, Des Marais DL, McKay J et al. (2012) Characterizing genomic variation of Arabidopsis thaliana: the roles of geography and climate. Molecular Ecology, 21, 5512–5529.

Laugen AT, Kruuk LEB, Laurila A et al. (2005a) Quantitative genetics of larval life-history traits in Rana temporaria in different environmental conditions. Genetics Research, 86, 161–170.

Laugen AT, Laurila A, Jönsson KI, Söderman F, Merilä J (2005b) Do common frogs (*Rana temporaria*) follow Bergmann’s rule? Evolutionary Ecology Research, 7, 717–731.

Laugen AT, Laurila A, Merilä J (2002) Maternal and genetic contributions to geographical variation in Rana temporaria larval life-history traits. Biological Journal fo the Linnean Society, 76, 61–70.

Laugen AT, Laurila A, Merilä J (2003a) Latitudinal and temperature-dependent variation in embryonic development and growth in Rana temporaria. Oecologia, 135, 548–554.

Laugen AT, Laurila A, Rasanen K, Merilä J (2003b) Latitudinal countergradient variation in the common frog (Rana temporaria) development rates – evidence for local adaptation. Journal of Evolutionary Biology, 16, 996–1005.

Laurila A, Karttunen S, Meril J (2002) Adaptive Phenotypic Plasticity and Genetics of Larval Life Histories in Two Rana Temporaria Populations. Evolution, 56, 617–627.

Laurila A, Pakkasmaa S, Merila J (2001) Influence of Seasonal Time Constraints on Growth and Development of Common Frog Tadpoles: A Photoperiod Experiment. Oikos, 95, 451–460.

Legendre P, Fortin M-J (2010) Comparison of the Mantel test and alternative approaches for detecting complex multivariate relationships in the spatial analysis of genetic data. Molecular Ecology Resources, 10, 831–844.

Legendre P, Legendre LF (2012) Numerical ecology. Elsevier.

Librado P, Rozas J (2009) DnaSP v5: a software for comprehensive analysis of DNA polymorphism data. Bioinformatics Application Note, 25, 1451–1452.

Lindgren B, Laurila A (2005) Proximate causes of adaptive growth rates: growth efficiency variation among latitudinal populations of Rana temporaria. Journal of Evolutionary Biology, 18, 820–828.

Lindgren B, Laurila A (2009) Physiological variation along a geographical gradient: is growth rate correlated with routine metabolic rate in Rana temporaria tadpoles? Biological Journal of the Linnean Society, 98, 217–224.

Loman J, Claesson D (2003) Plastic response to pond drying in tadpoles Rana temporaria: tests of cost models. Evolutionary Ecology Research, 5, 179–194.

Lugon-Moulin N, Hausser J (2002) Phylogeographical structure, postglacial recolonisation and barriers to gene flow in the distinctive Valais chromosome race of the common shrew (Sorex araneus). Molecular Ecology, 11, 785–794.

Luikart G, England PR, Tallmon D, Jordan S, Taberlet P (2003) The power and promise of population genomics: from genotyping to genome typing. Nature Reviews Genetics, 4, 981–94.

Luu K, Bazin E, Blum MGB (2016) pcadapt: An R package to perform genome scans for selection based on principal component analysis. Molecular Ecology Resources, 33, 67–77.

Manel S, Perrier C, Pratlong M et al. (2016) Genomic resources and their influence on the detection of the signal of positive selection in genome scans. Molecular ecology, 25, 170–184.

Manel S, Poncet BN, Legendre P, Gugerli F, Holderegger R (2010) Common factors drive adaptive genetic variation at different spatial scales in Arabis alpina. Molecular ecology, 19, 3824–35.

Marquis O, Miaud C (2008) Variation in UV sensitivity among common frog Rana temporaria populations along an altitudinal gradient. Zoology, 111, 309–317.

Marquis O, Miaud C, Lena J-P (2008) Developmental responses to UV-B radiation in common frog Rana temporaria embryos from along an altitudinal gradient. Population Ecology, 50, 123–130.

Mátyás G, Sperisen C (2001) Chloroplast DNA polymorphisms provide evidence for postglacial re-colonisation of oaks (Quercus spp.) across the Swiss Alps. Theoretical and Applied Genetics, 102, 12–20.

McCain CM, Colwell RK (2011) Assessing the threat to montane biodiversity from discordant shifts in temperature and precipitation in a changing climate. Ecology Letters, 14, 1236–1245.

Merila J, Laurila A, Laugen AT, Rasanen K, Pahkala M (2000) Plasticity in age and size at metamorphosis in Rana temporaria: comparison of high and low latitude populations. Ecography, 23, 457–465.

Merilä J, Laurila A, Laugen AT, Räsänen K, Pahkala M (2000) Plasticity in Age and Size at Metamorphosis in Rana Temporaria: Comparison of High and Low Latitude Populations. Ecography, 23, 457–465.

Messer PW, Petrov DA (2013) Population genomics of rapid adaptation by soft selective sweeps. Trends in Ecology and Evolution, 28, 659–669.

Miaud C, Guyetant R, Elmberg J (1999) Variations in life-history traits in the common frog Rana temporaria (Amphibia: Anura): a literature review and new data from the French Alps. Journal of Zoology, 249, 61–73.

Miaud C, Merilä J (2001) Local adaptation or environmental induction? Causes of population differentiation in alpine amphibians. Biota, 2, 31–50.

Muir AP, Biek R, Mable BK (2014a) Behavioural and physiological adaptations to low-temperature environments in the common frog, Rana temporaria. BMC Evolutionary Biology, 14, 1471–2148.

Muir A, Piek R, Thomas R, Mable B (2014b) Local adaptation with high gene flow: temperature parameters drive adaptation to altitude in the common frog (Rana temporaria). Molecular ecology, 23, 561–574.

Nychka D, Furrer R, Paige J, Sain S (2015) fields: Tools for spatial data.

Oksanen J, Blanchet FG, Kindt R et al. (2015) Vegan: community ecology package. R package vegan, vers. 2.2-1.

Pahkala M, Laurila A, Meril J (2002) Effects of ultraviolet-B radiation on common frog Rana temporaria embryos from along a latitudinal gradient. Oecologia, 133, 458–465.

Pahkala M, Laurila A, Merila J (2000) Ambient Ultraviolet-B radiation reduces hatchling size in the common frog Rana temporaria. Ecography, 23, 531–538.

Pahkala M, Laurila A, Merila J (2001) Carry-over effects of ultraviolet-B radiation on larval tness in Rana temporaria. Proceedings of the Royal Society of London B: Biological Sciences, 268, 1699–1706.

Pahkala M, Merila J, Ots I, Laurila A (2003) Effects of ultraviolet-B radiation on metamorphic traits in the common frog Rana temporaria. Journal of the Zoological Society of London, 259, 57–62.

Palo JU, O’Hara RB, Laugen AT et al. (2003a) Latitudinal divergence of common frog (Rana temporaria) life history traits by natural selection: evidence from a comparison of molecular and quantitative genetic data. Molecular ecology, 12, 1963–1978.

Palo JU, O’Hara RB, Laugen AT et al. (2003b) Latitudinal divergence of the common frog (Rana temporaria) life history traits by natural selection: evidence from a comparison of molecular quantitative genetic data. Molecular Ecology, 12, 1963–1978.

Palo JU, Schmeller DS, Laurila A et al. (2004) High degree of population subdivision in a widespread amphibian. Molecular Ecology, 13, 2631–2644.

Parisod C (2008) Postglacial recolonisation of plants in the western Alps of Switzerland. Botanica Helvetica, 118, 1–12.

Parmesan C (2006) Ecological and Evolutionary Responses to Recent Climate Change. Annual Review of Ecology, Evolution, and Systematics, 37, 637–669.

Parmesan C, Yohe G (2003) A globally coherent fingerprint of climate change impacts across natural systems. Nature, 421, 37–42.

Pennings PS, Hermisson J (2006) Soft sweeps II - Molecular population genetics of adaptation from recurrent mutation or migration. Molecular Biology and Evolution, 23, 1076–1084.

Peterson BK, Weber JN, Kay EH, Fisher HS, Hoekstra HE (2012) Double Digest RADseq: An Inexpensive Method for De Novo SNP Discovery and Genotyping in Model and Non-Model Species. PloS one, 7.

Polechová J, Barton NH (2015) Limits to adaptation along environmental gradients. Proceedings of the National Academy of Sciences of the United States of America, 112, 6401–6406.

Poncet BN, Herrmann D, Gugerli F et al. (2010) Tracking genes of ecological relevance using a genome scan in two independent regional population samples of Arabis alpina. Molecular ecology, 19, 2896–907.

Pritchard JK, Di Rienzo A (2010) Adaptation - not by sweeps alone. Nature Reviews Genetics, 11, 665–7.

Purcell S, Neale B, Todd-Brown K et al. (2007) PLINK: A tool set for whole-genome association and population-based linkage analyses. American Journal of Human Genetics, 81, 559–575.

Raj A, Stephens M, Pritchard JK (2014) FastSTRUCTURE: Variational inference of population structure in large SNP data sets. Genetics, 197, 573–589.

Rehm EM, Olivas P, Stroud J, Feeley KJ (2015) Losing your edge: climate change and the conservation value of range-edge populations. Ecology and evolution, 5, 4315–4326.

Rellstab C, Fischer MC, Zoller S et al. (2016) Local adaptation (mostly) remains local: reassessing environmental associations of climate-related candidate SNPs in Arabidopsis halleri. Heredity, 1–9.

Rellstab C, Gugerli F, Eckert AJ, Hancock AM, Holderegger R (2015) A practical guide to environmental association analysis in landscape genomics. Molecular Ecology, 24, 4348–70.

Rigling A, Bigler C, Eilmann B et al. (2013) Driving factors of a vegetation shift from Scots pine to pubescent oak in dry Alpine forests. Global Change Biology, 19, 229–240.

Rodrigues N, Betto-Colliard C, Jourdan-Pineau H, Perrin N (2013) Within-population polymorphism of sex-determination systems in the common frog (Rana temporaria). Journal of evolutionary biology, 26, 1569–1577.

Roesti M, Salzburger W, Berner D (2012) Uninformative polymorphisms bias genome scans for signatures of selection. BMC Evolutionary Biology, 12, 94.

Roff DA (1996) The evolution of threshold traits in animals. The Quarterly Review of Biology, 71, 3–35.

Rogers AR, Harpending H (1992) Population Growth Makes Waves in the Distribution of Pairwise Genetic Differences. Molecular biology and evolution, 9, 552–569.

Rogivue A, Graf R, Parisod C, Holderegger R, Gugerli F (2018) The phylogeographic structure of Arabis alpina in the Alps shows consistent patterns across different types of molecular markers and geographic scales. Alpine Botany, 0, 0.

Rousset F (1997) Genetic differentiation and estimation of gene flow from F-statistics under isolation by distance. Genetics, 145, 1219–1228.

Roy K, Valentine JW, Jablonski D, Kidwell SM (1996) Scales of climatic variability and time averaging in Pleistocene biotas: implications for ecology and evolution. Trends in Ecology & Evolution, 11, 458–463.

Schmeller DS, Palo JU, Merilä J (2008) A contact zone between two distinct Rana temporaria lineages in northern Germany. Alytes, 25, 93–98.

Schneider S, Excoffier L (1999) Estimation of Past Demographic Parameters From the Distribution of Pairwise Differences When the Mutation Rates Vary Among Sites: Application to Human Mitochondrial DNA. Genetics, 152, 1079–1089.

Schweizer RM, VonHoldt BM, Harrigan R et al. (2016) Genetic subdivision and candidate genes under selection in North American grey wolves. Molecular Ecology, 25, 380–402.

Sillero N, Campos J, Bonardi A et al. (2014) Updated distribution and biogeography of amphibians and reptiles of Europe. Amphibia-Reptilia, 35, 1–31.

Sork VL, Aitken SN, Dyer RJ et al. (2013) Putting the landscape into the genomics of trees: approaches for understanding local adaptation and population responses to changing climate. Tree Genetics & Genomes, 9, 901–911.

Ståhlberg F, Olsson M, Uller T (2001) Population divergence of developmental thermal optima in Swedish common frogs, Rana temporaria. Journal of Evolutionary Biology, 14, 755–762.

Stamatakis A (2006) RAxML-VI-HPC: maximum likelihood-based phylogenetic analyses with thousands of taxa and mixed models. Bioinformatics Application Note, 22, 2688–2690.

Stapley J, Reger J, Feulner PGD et al. (2010) Adaptation genomics: the next generation. Trends in ecology & evolution, 25, 705–12.

Stefani F, Gentilli A, Sacchi R et al. (2012) Refugia within refugia as a key to disentangle the genetic pattern of a highly variable species: The case of Rana temporaria Linnaeus, 1758 (Anura, Ranidae). Molecular Phylogenetics and Evolution, 65, 718–726.

Stehlik I, Blattner FR, Holderegger R, Bachmann K (2002) Nunatak survival of the high Alpine plant Eritrichium nanum (L.) Gaudin in the central Alps during the ice ages. Molecular Ecology, 11, 2027–2036.

Taberlet P, Fumagalli L, Wust-Saucy A-G, Cossons J-F (1998) Comparative phylogeography and postglacial colonization routes in Europe. Molecular Ecolgy, 7, 453–464.

Teacher AGF, Garner TWJ, Nichols RA (2009) European phylogeography of the common frog (Rana temporaria): routes of postglacial colonization into the British Isles, and evidence for an Irish glacial refugium. Heredity, 102, 490–6.

Templeton AR, Crandall KA, Sing CF (1992) Cladistic Analysis of Phenotypic Associations With Haplotypes Inferred From Restriction Endonuclease Mapping and DNA Sequence Data. III. Cladogram Estimation. Genetics, 132, 619–633.

Thomassen H a, Cheviron Z a, Freedman AH et al. (2010) Spatial modelling and landscape-level approaches for visualizing intra-specific variation. Molecular Ecology, 19, 3532–48.

Tinner W, Theurillat J (2003) Uppermost limit, extent, and fluctuations of the timberline and treeline ecocline in the Swiss Central Alps during the past 11,500 years. Arctic, Antarctic, and Alpine Research, 35, 158–169.

Veith M, Kosuch J, Vences M (2003) Climatic oscillations triggered post-Messinian speciation of Western Palearctic brown frogs (Amphibia, Ranidae). Molecular phylogenetics and evolution, 26, 310–327.

Veith M, Vences M, Vieites DR, Nieto-roman S, Palanca A (2002) Genetic differentiation and population structure within Spanish common frogs (Rana temporaria complex; Ranidae, Amphibia). Folia Zoologica, 51, 307–318.

Vences M, Hauswaldt JS, Steinfartz S et al. (2013) Radically different phylogeographies and patterns of genetic variation in two European brown frogs, genus Rana. Molecular phylogenetics and evolution, 68, 657–70.

Vergeer P, Kunin WE (2013) Adaptation at range margins: common garden trials and the performance of Arabidopsis lyrata across its northwestern European range. New Phytologist, 197, 989–1001.

Vitti JJ, Grossman SR, Sabeti PC (2013) Detecting Natural Selection in Genomic Data. Annu. Rev. Genet, 47, 97–120.

Wang IJ (2012) Environmental and topographic variables shape genetic structure and effective population sizes in the endagered Yosemite toad. Diversity and Distributions, 18, 1033–1041.

Wang IJ, Glor RE, Losos JB (2013) Quantifying the roles of ecology and geography in spatial genetic divergence. Ecology letters, 16, 175–82.

Weir BS, Cockerham CC (1984) Estimating F-Statistics for the Analysis of Population Structure. Evolution, 38, 1358–1370.

Wigginton JE, Cutler DJ, Abecasis GR (2005) A note on exact tests of Hardy-Weinberg equilibrium. American Journal of Human Genetics, 76, 887–93.

Willis KJ, Whittaker RJ (2008) The refugial debate. Science, 287, 1406–1407.

Yannic G, Basset P, Hausser J (2008) Phylogeography and recolonisation of the Swiss Alps by the Valais shrew (Sorex antinorii), inferred with autosomal and sex-specific markers. Molecular Ecology, 17, 4118–4133.

Yannic G, Pellissier L, Dubey S et al. (2012) Multiple refugia and barriers explain the phylogeography of the Valais shrew, Sorex antinorii (Mammalia: Soricomorpha). Biological Journal of the Linnean Society, 105, 864–880.

Yeaman S (2015) Local Adaptation by Alleles of Small Effect. The American Naturalist, 186, S74–S89.

