## Supplemental File for "European common frog (*Rana temporaria*) recolonised Switzerland from multiple glacial refugia in northern Italy via trans- and circum-Alpine routes"

***Supplemental Material***

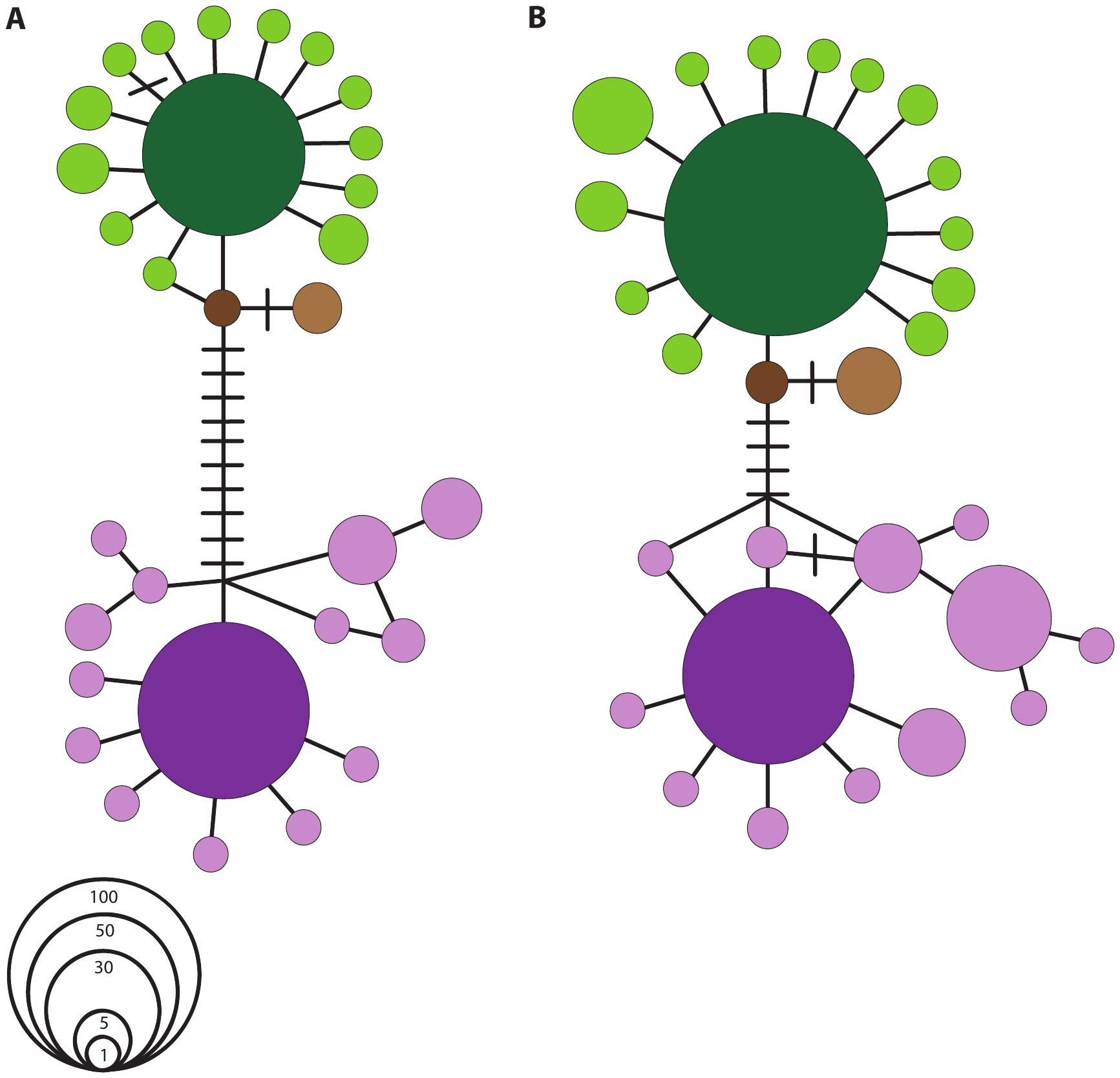

***Figure S1*** Haplotype network of individuals sampled in Switzerland only. a) 448bp cytb sequenced for 368 individuals from 72 populations, b) 1076bp of mitochondrial DNA (448bp cytb and 628bp COX1) sequenced for 44 populations (151 individuals) from Switzerland. Circles represent haplotypes, and are scaled by sample size. Colours correspond to those in Fig. 1. A solid line represents a single mutational step linking two haplotypes, with cross hatches added for additional steps.

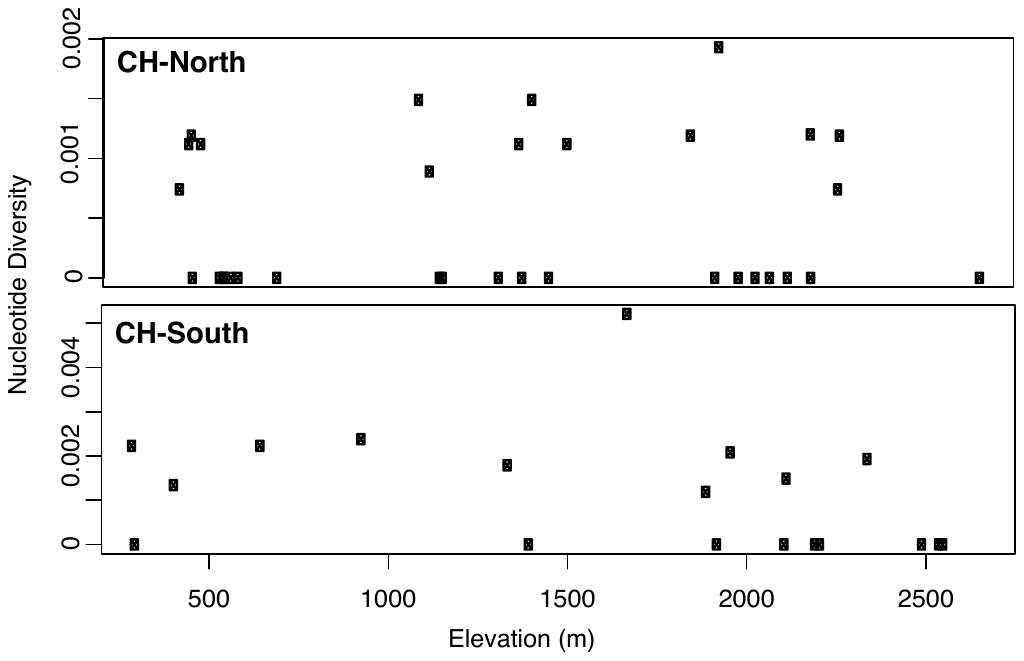

***Figure S2*** Nucleotide diversity plotted across elevation for CH-North (top) and CH-South (bottom) from Switzerland. We excluded populations where the haplogroups co-occur.

***Table S1*** Populations sampled and sequenced at cytochrome b. N= number of samples sequenced, Haplotypes= number of haplotypes found in the population, Hd (SD)= Haplotype diversity (standard deviation), nd (SD) = nucleotide diversity (standard deviation).

| **Population** | **Lat.** | **Long.** | **Elevation** | **N** | **length (bp)** | **Haplotypes** | **Hd (SD)** | **nd (SD)** |
| --- | --- | --- | --- | --- | --- | --- | --- | --- |
| agra | 46.03 | 8.90 | 930 | 6 | 448 | 3 | 0.8 (0.122) | 0.00238 (0.0005) |
| alpl | 46.94 | 8.66 | 1506 | 4 | 448 | 2 | 0.5 (0.27) | 0.00112 (0.00059) |
| apla | 46.79 | 9.50 | 2118 | 6 | 448 | 3 | 0.6 (0.22) | 0.0091 (0.00297) |
| arce | 46.16 | 8.75 | 408 | 6 | 448 | 3 | 0.73 (0.16) | 0.0064 (0.00251) |
| bach | 46.67 | 8.03 | 2265 | 6 | 448 | 2 | 0.53 (0.17) | 0.00119 (0.00038) |
| bela | 46.38 | 7.98 | 2179 | 6 | 448 | 3 | 0.73 (0.16) | 0.00699 (0.00288) |
| bend | 46.49 | 6.88 | 590 | 6 | 448 | 1 | 0 (0) | 0 (0) |
| berm | 47.02 | 7.52 | 550 | 6 | 448 | 1 | 0 (0) | 0 (0) |
| bide | 47.14 | 9.40 | 1983 | 6 | 448 | 1 | 0 (0) | 0 (0) |
| birk | 47.30 | 8.81 | 550 | 4 | 448 | 1 | 0 (0) | 0 (0) |
| bnnp | 46.41 | 10.03 | 2342 | 6 | 448 | 3 | 0.73 (0.16) | 0.00193 (0.00055) |
| buel | 46.86 | 9.73 | 2260 | 6 | 448 | 2 | 0.33 (0.22) | 0.00074 (0.00048) |
| cava | 46.36 | 9.03 | 2003 | 6 | 448 | 2 | 0.33 (0.22) | 0.00521 (0.00336) |
| csan | 46.32 | 7.30 | 2110 | 6 | 448 | 1 | 0 (0) | 0 (0) |
| egel | 47.25 | 9.50 | 445 | 6 | 448 | 2 | 0.33 (0.22) | 0.00595 (0.00384) |
| fada | 46.98 | 9.60 | 1093 | 3 | 448 | 2 | 0.67 (0.31) | 0.00149 (0.00070) |
| fess | 47.02 | 9.14 | 2184 | 4 | 448 | 2 | 0.5 (0.27) | 0.0012 (0.00059) |
| flue | 46.75 | 9.95 | 2388 | 3 | 448 | 3 | 0.83 (0.05) | 0.0115 (0.00335) |
| forn | 46.48 | 8.58 | 2089 | 5 | 448 | 3 | 0.7 (0.22) | 0.00982 (0.00295) |
| full | 46.17 | 7.10 | 2074 | 5 | 448 | 3 | 0.7 (0.22) | 0.00759 (0.00333) |
| fuor | 46.44 | 9.83 | 2494 | 6 | 448 | 1 | 0 (0) | 0 (0) |
| gdwe | 46.64 | 8.07 | 1380 | 6 | 448 | 1 | 0 (0) | 0 (0) |
| gola | 46.10 | 8.97 | 975 | 6 | 448 | 5 | 0.93 (0.12) | 0.00774 (0.00282) |
| gott | 46.59 | 8.56 | 2116 | 6 | 448 | 2 | 0.33 (0.22) | 0.00149 (0.00096) |
| grma | 46.24 | 7.01 | 425 | 6 | 448 | 3 | 0.6 (0.22) | 0.00714 (0.00356) |
| grsh | 46.66 | 8.10 | 1929 | 6 | 448 | 3 | 0.73 (0.16) | 0.00193 (0.00055) |
| gruu | 46.86 | 9.79 | 2120 | 4 | 448 | 1 | 0 (0) | 0 (0) |
| hdns | 46.85 | 9.76 | 1918 | 4 | 448 | 1 | 0 (0) | 0 (0) |
| jagg | 46.74 | 8.06 | 570 | 4 | 448 | 1 | 0 (0) | 0 (0) |
| kand | 46.63 | 7.69 | 698 | 4 | 448 | 1 | 0 (0) | 0 (0) |
| kebe | 47.54 | 8.78 | 453 | 4 | 448 | 2 | 0.5 (0.27) | 0.00112 (0.00059) |
| lens | 46.29 | 7.46 | 1338 | 5 | 448 | 2 | 0.4 (0.24) | 0.00179 (0.00106) |
| lucm | 46.56 | 8.80 | 1922 | 6 | 448 | 1 | 0 (0) | 0 (0) |
| magn | 46.43 | 8.68 | 1840 | 4 | 448 | 2 | 0.5 (0.27) | 0.0067 (0.00355) |
| mart | 46.03 | 8.94 | 407 | 5 | 448 | 2 | 0.6 (0.18) | 0.00134 (0.00039) |
| mctn | 46.48 | 9.72 | 2542 | 5 | 448 | 1 | 0 (0) | 0 (0) |
| mgns | 46.25 | 6.85 | 1372 | 4 | 448 | 2 | 0.5 (0.27) | 0.00112 (0.00059) |
| moau | 46.34 | 6.79 | 1850 | 6 | 448 | 2 | 0.53 (0.17) | 0.00119 (0.00038) |
| moir | 46.10 | 7.57 | 2553 | 4 | 448 | 1 | 0 (0) | 0 (0) |
| muet | 47.45 | 8.61 | 460 | 6 | 448 | 2 | 0.53 (0.17) | 0.00119 (0.00038) |
| munt | 46.73 | 9.44 | 648 | 2 | 448 | 2 | 1 (0.5) | 0.00223 (0.00112) |
| oalp | 46.65 | 8.64 | 1960 | 6 | 448 | 3 | 0.73 (0.16) | 0.00208 (0.00058) |
| otte | 46.73 | 7.36 | 1455 | 6 | 448 | 1 | 0 (0) | 0 (0) |
| petl | 45.89 | 7.15 | 2655 | 4 | 448 | 1 | 0 (0) | 0 (0) |
| pizo | 46.98 | 9.42 | 2209 | 4 | 448 | 1 | 0 (0) | 0 (0) |
| pozz | 46.35 | 8.96 | 290 | 4 | 448 | 3 | 0.83 (0.22) | 0.00223 (0.00076) |
| prad | 46.78 | 9.53 | 1449 | 5 | 448 | 4 | 0.9 (0.16) | 0.01161 (0.00307) |
| roes | 46.90 | 7.20 | 538 | 6 | 448 | 1 | 0 (0) | 0 (0) |
| rose | 46.14 | 7.05 | 450 | 4 | 448 | 3 | 0.83 (0.22) | 0.01042 (0.00367) |
| rotc | 47.01 | 9.31 | 2185 | 4 | 448 | 1 | 0 (0) | 0 (0) |
| rusc | 46.37 | 7.24 | 1315 | 4 | 448 | 1 | 0 (0) | 0 (0) |
| sali | 46.26 | 8.69 | 331 | 4 | 448 | 3 | 0.83 (0.22) | 0.00967 (0.00383) |
| saxm | 47.07 | 9.38 | 463 | 9 | 448 | 1 | 0 (0) | 0 (0) |
| scai | 45.98 | 8.93 | 298 | 2 | 448 | 1 | 0 (0) | 0 (0) |
| seeo | 47.05 | 9.58 | 2030 | 6 | 448 | 1 | 0 (0) | 0 (0) |
| seji | 46.80 | 9.73 | 2090 | 6 | 448 | 2 | 0.83 (0.22) | 0.00595 (0.00384) |
| shwe | 47.19 | 9.33 | 1159 | 6 | 448 | 1 | 0 (0) | 0 (0) |
| siec | 46.99 | 9.55 | 517 | 5 | 448 | 2 | 0.6 (0.18) | 0.00938 (0.00274) |
| star | 46.27 | 8.77 | 1892 | 6 | 448 | 2 | 0.53 (0.17) | 0.00119 (0.00038) |
| stba | 46.49 | 9.17 | 2059 | 6 | 448 | 2 | 0.33 (0.22) | 0.00521 (0.00336) |
| stir | 46.53 | 7.55 | 2070 | 4 | 448 | 1 | 0 (0) | 0 (0) |
| stls | 46.97 | 9.75 | 1672 | 6 | 448 | 2 | 0.33 (0.22) | 0.00521 (0.00336) |
| tana | 46.35 | 6.84 | 1408 | 6 | 448 | 3 | 0.6 (0.22) | 0.00149 (0.00062) |
| trim | 46.90 | 9.54 | 545 | 5 | 448 | 2 | 0.4 (0.24) | 0.00625 (0.00371) |
| trpa | 46.29 | 7.28 | 2196 | 4 | 448 | 1 | 0 (0) | 0 (0) |
| tsee | 46.55 | 7.75 | 1151 | 6 | 448 | 1 | 0 (0) | 0 (0) |
| vilt | 47.03 | 9.45 | 486 | 4 | 448 | 2 | 0.5 (0.27) | 0.00112 (0.00059) |
| vora | 47.16 | 9.38 | 1123 | 5 | 448 | 2 | 0.4 (0.24) | 0.00089 (0.00053) |
| wale | 47.12 | 9.10 | 427 | 6 | 448 | 2 | 0.33 (0.22) | 0.00074 (0.00048) |
| wdji | 46.81 | 9.72 | 1638 | 6 | 448 | 3 | 0.8 (0.12) | 0.01071 (0.00303) |
| wise | 47.19 | 9.48 | 445 | 5 | 448 | 4 | 0.9 (0.16) | 0.01027 (0.00247) |
| zeni | 46.52 | 8.89 | 1397 | 5 | 448 | 1 | 0 (0) | 0 (0) |

***Table S2*** Populations sampled and sequenced at COX1. N = number of samples sequenced, Haplotypes = number of haplotypes found in the population, Hd (SD) = Haplotype diversity (standard deviation), nd (SD) = nucleotide diversity (standard deviation).

| **Population** | **N** | **length (bp)** | **nr haplotypes** | **Hd (SD)** | **nucleotide diversity (SD)** |
| --- | --- | --- | --- | --- | --- |
| alpl | 4 | 628 | 1 | 0 (0) | 0 (0) |
| arce | 3 | 628 | 1 | 0 (0) | 0 (0) |
| bach | 4 | 628 | 1 | 0 (0) | 0 (0) |
| bela | 3 | 628 | 1 | 0 (0) | 0 (0) |
| bide | 3 | 628 | 2 | 0.67 (0.31) | 0.00106 (0.00050) |
| birk | 4 | 628 | 1 | 0 (0) | 0 (0) |
| bnnp | 3 | 628 | 2 | 0.67 (0.31) | 0.00106 (0.00050) |
| cava | 3 | 628 | 2 | 0.67 (0.31) | 0.00637 (0.003) |
| egel | 3 | 628 | 2 | 0.67 (0.31) | 0.00106 (0.00050) |
| fada | 3 | 628 | 1 | 0 (0) | 0 (0) |
| flue | 4 | 628 | 3 | 0.83 (0.22) | 0.00717 (0.00207) |
| forn | 4 | 628 | 3 | 0.83 (0.22) | 0.00717 (0.00207) |
| fuor | 3 | 628 | 1 | 0 (0) | 0 (0) |
| gott | 4 | 628 | 2 | 0.5 (0.27) | 0.0008 (0.00042) |
| grma | 4 | 628 | 2 | 0.5 (0.27) | 0.0008 (0.00042) |
| grsh | 4 | 628 | 1 | 0 (0) | 0 (0) |
| gruu | 4 | 628 | 2 | 0.5 (0.27) | 0.0008 (0.00042) |
| hdns | 4 | 628 | 2 | 0.5 (0.27) | 0.0008 (0.00042) |
| jagg | 4 | 628 | 1 | 0 (0) | 0 (0) |
| kand | 4 | 628 | 2 | 0.5 (0.27) | 0.00159 (0.00084) |
| kebe | 4 | 628 | 1 | 0 (0) | 0 (0) |
| magn | 3 | 628 | 2 | 0.67 (0.31) | 0.00743 (0.0035) |
| mart | 4 | 628 | 2 | 0.67 (0.2) | 0.00212 (0.00065) |
| mctn | 4 | 628 | 1 | 0 (0) | 0 (0) |
| mgns | 4 | 628 | 1 | 0 (0) | 0 (0) |
| moir | 4 | 628 | 1 | 0 (0) | 0 (0) |
| petl | 4 | 628 | 1 | 0 (0) | 0 (0) |
| pizo | 4 | 628 | 1 | 0 (0) | 0 (0) |
| pozz | 4 | 628 | 2 | 0.5 (0.27) | 0.00159 (0.00084) |
| prad | 4 | 628 | 2 | 0.5 (0.27) | 0.00478 (0.00253) |
| rotc | 4 | 628 | 1 | 0 (0) | 0 (0) |
| rusc | 4 | 628 | 1 | 0 (0) | 0 (0) |
| seeo | 4 | 628 | 2 | 0.67 (0.2) | 0.00106 (0.00033) |
| shwe | 4 | 628 | 1 | 0 (0) | 0 (0) |
| siec | 1 | 628 | 1 | 0 (0) | 0 (0) |
| star | 1 | 628 | 1 | 0 (0) | 0 (0) |
| stba | 3 | 628 | 3 | 1.0 (0.27) | 0.00743 (0.00304) |
| stir | 2 | 628 | 1 | 0 (0) | 0 (0) |
| stls | 1 | 628 | 1 | 0 (0) | 0 (0) |
| tana | 3 | 628 | 1 | 0 (0) | 0 (0) |
| trpa | 4 | 628 | 1 | 0 (0) | 0 (0) |
| vora | 2 | 628 | 1 | 0 (0) | 0 (0) |
| wdji | 4 | 628 | 2 | 0.5 (0.27) | 0.00478 (0.00253) |
| zeni | 3 | 628 | 1 | 0 (0) | 0 (0) |
